# Prosperity of the commons: Generalist mycorrhizal species dominate a mixed forest and may promote forest diversity by mediating resource sharing among trees

**DOI:** 10.1101/2022.08.01.502298

**Authors:** Ido Rog, Ohad Lewin-Epstein, Stav Livne-Luzon, Lilach Hadany, Tamir Klein

**Author notes:** These authors contributed equally to this study (Ido Rog, Ohad Lewin-Epstein). Corresponding author: Tamir Klein,.

## Abstract

Mechanisms of host–microbe interactions and their direct impact on both parties have been extensively researched, however, much less is known on the effect of these interactions on the ecology of the host-community. Here we investigate tree-fungi mycorrhizal interactions, focusing on mycorrhizal-mediated resource sharing among trees, while examining the dynamics between specialist and generalist fungi and their implications on the forest ecology. Using genetic meta-barcoding, we identified the fungal community colonizing different trees in a mixed forest, and generated an extensive mapping connecting fungal sequences to their tree hosts. The mycorrhizal fungal community diverged between ectomycorrhizal and arbuscular host trees, but, unexpectedly, multiple ectomycorrhizal species colonized roots of non-ectomycorrhizal host trees. We complemented these findings by a novel computational framework, modeling competition between generalist and specialist mycorrhizal fungi, accounting for fungal-mediated resource sharing among neighboring trees. The analysis of the model revealed that generalist mycorrhizal networks may affect the entire tree community, and contribute to the maintenance of forest diversity in the long run. Furthermore, higher initial spatial mixing of trees can promote the evolution of generalist mycorrhizal species. These novel belowground interactions among trees and fungi may significantly impact forest biodiversity.

## Introduction

The symbiosis between microbes and their hosts strongly influences the microbial community as well as the host wellbeing and functioning (Zilber-Rosenberg and Rosenberg 2008, Kinross et al. 2011, Chaparro et al. 2014, Goodrich et al. 2014, Sasse et al. 2018, Trivedi et al. 2020). Microbial symbionts vary in their degree of host specificity (Bruns et al. 2002, Bäumler and Fang 2013, Semchenko et al. 2022), defined as the rate at which a specific microbial species associates with a particular host species, while having lower symbiotic benefits with alternative hosts (Kiers et al. 2011) across a large geographical range (Wolfe and Pringle 2012). Different levels of specialism may differently project on the host-microbe interactions and their impact on the diversity and structure of the host community, and thus may play a crucial ecological role in shaping microbial communities. While mechanisms of host–microbe interactions and their direct impact on both parties have been extensively researched, less is known on how host-microbe interactions can cascade up to affect the ecology of the entire community. The symbiosis between soil microbes and tree roots has long been studied, revealing how strongly it influences the functioning of the tree and the residing microbial community, both below and above the ground (Wardle et al. 2004). However, the impact of this symbiosis on the forest configuration and specifically on forest heterogeneity and spatial structure are less known.

In natural environments, soil nutrients are usually limited, and most plants rely on symbiotic fungi for nutrient uptake. Fungi form symbiotic interactions with the plant roots (Mycorrhiza) based on the exchange of host-derived C products and soil-derived mineral nutrients, in contrast to saprotrophic fungi that depend on decayed organic matter as their sole C source. Mycorrhizal fungi have been shown to uptake soil nutrients, improve plants’ functioning, and are the dominant fungi associated with tree roots in many habitats (Nara and Hogetsu 2004, Smith and Read 2010). The two most common types of mycorrhizal associations characterizing plant communities are Ectomycorrhizal fungi (EMF) and Arbuscular fungi (AMF) (Smith and Read 2010). While AMF have been shown to play a crucial role in phosphorus uptake, EMF have specialized in habitats poor of nitrogen. However, both of these groups are distributed all around the globe and their distribution is strongly influenced by the climate of certain biomes (Read 1991, Steidinger et al. 2019). For example, EMF host trees are located mostly in seasonally cold and dry climates, while AMF host trees prevail in seasonally warm and wet climates. Globally, EMF plant species are more conservative in their use of nitrogen (and sometimes, phosphorous too) than AMF plant species (Averill et al. 2019). Mixed forests that are dominated by both AMF and EMF host trees and prevail in many areas around the globe, provide a unique opportunity to assess what happens when these different guilds of hosts and symbionts meet.

Most mycorrhizal fungal communities are characterized with a few dominant species and a long ‘tail’ of less abundant, rare, species. One of the theories suggested to explain this mycorrhizal community characteristic relies on the ability of most mycorrhizal fungal species (Kennedy 2010), specifically EMF, to colonize multiple occurring hosts at the same time (Molina 1992) while expressing various host preferences (Ishida et al. 2007, Tedersoo et al. 2008, Smith et al. 2009, Pickles et al. 2012, Molina and Horton 2015). Various compatibility levels between the host and the fungi can lead to mycorrhizal colonization of multiple tree genera (“Generalists”) where as other mycorrhizal fungi that colonize mainly one tree genus (“Specialists”) (Horton and Bruns 1998, Wolfe and Pringle 2012). Mycorrhizal fungal species exhibiting low host specificity are usually among the most abundant mycorrhizal taxa in specific plant communities (Horton and Bruns 1998). The degree of host-fungi resource exchange can vary substantially, and one plant can benefit more from being connected to mycorrhizal fungi than other plants (Van Der Heijden and Horton 2009). For example, *Linum ustiatissimum* allocates small amounts of C to its mycorrhizal fungi but receives up to 90% of the plant demands for nitrogen and phosphorus from its mycorrhizal symbionts (Walder et al. 2012). In contrast, *Sorghum bicolor* allocates most of the assimilated C to its mycorrhizal fungi but takes only a small amount of nutrients in return. Colonizing multiple occurring hosts with different resource exchange properties (i.e. amount, chemical compound and, dynamic of C) could be highly beneficial to the fungi, especially in the mixed forest characterized by high diversity of hosts occupying various niches.

Multiple studies revealed an additional facet of the tree-fungi mycorrhizal interactions, portrayed in the ability of mycorrhizal fungi to mediate resource sharing among trees, by transporting C via the mycorrhizal hyphae that is connected to neighboring trees (Simard et al. 1997, Walder and van der Heijden 2015, Klein et al. 2016). A single fungal genet can be associated with many root tips of different trees to form belowground links among trees, facilitating the transfer of C, nutrients, and water from one tree to another. These Mycorrhizal hyphae can share C between trees of different ages (Teste et al. 2009), different species (Simard et al. 1997), and even between conifers and broadleaves trees (Klein et al. 2016). However, not all trees are connected, and various guilds are formed in each specific forest (Pickles and Simard 2017). The amount and direction of C transfer in the mycorrhizal networks have been shown to be controlled by phylogenetic relatedness (Rog et al. 2020), kin association (Pickles et al. 2017), and other parameters. In Mediterranean saplings, it was calculated that 0.5% of the root C was assimilated in the neighbor tree (Cahanovitc et al. 2022), while in the temperate mature forest; it reached to even 4% of the net forest assimilates (Klein et al. 2016). The C transfer dynamic through the mycorrhizal hyphae is still under investigation, but data from pulse labeling experiments present around ~85 cm year^-1^ with a maximum of 6 m in mature trees. These belowground interactions can potentially influence forest connectivity and resistance to future climate changes by redistribution of C among the various forest players.

The multiple evidence for fungal-mediated resource sharing among trees can be considered a form of cooperation between the trees, induced by mycorrhizal fungi. Mycorrhiza-induced resource sharing can hence serve as an example of the microbe-induced cooperation hypothesis (Lewin-Epstein et al. 2017, Gurevich et al. 2020, Lavy et al. 2020, Lewin-Epstein and Hadany 2020). According to this hypothesis, natural selection may favor microbes that increase the tendency of their hosts to cooperate with other hosts under a wide range of conditions, including when the hosts are not kin, and thus promote cooperative behavior in the host population. The two foundations of this hypothesis are the ability of microorganisms, to affect the behavior, wellbeing and functioning of their hosts (Poulin 2010, Sampson and Mazmanian 2015, Vandenkoornhuyse et al. 2015, Rosenberg and Zilber-Rosenberg 2018), and their ability to spread from one host to another via horizontal transmission. The integration of these two, facilitate a potential advantage for a microbial trait that mediates host-cooperation. Due to horizontal transmission, microbes that induce their host to cooperate with another host, could help their microbial kin that were transmitted to the recipient host - even when the hosts are not related.

Indeed, mycorrhizal fungi have been shown to impact their host wellbeing and mediate resource allocation between trees, and thus potentially induce cooperation (Gorzelak et al. 2015, Sapes et al. 2021), namely confer a cost to the donor and provide a benefit to its recipient. They also have the ability to spread horizontally, between neighboring trees, by expanding the mycelium and inhabiting new host trees (Nara and Hogetsu 2004). Following mycorrhizal fungi expansion from one tree to its neighbor, the two trees, will share closely related fungi. According to the theory, this combination may yield an evolutionary advantage to mycorrhizal fungal species that induce their tree hosts to cooperate and improve the wellbeing of their neighbors, which may host kin mycorrhizal fungi due to horizontal transmission. Central parameters for describing cooperation among hosts include its cost for the donor tree and its benefit for the recipient tree. In the case of resource sharing, the cost and the benefit can differ dramatically, for example, when carried out in asymmetric interactions (e.g., between young and mature host, large and small host, stressed and prosperous host, etc.), or when applied between different species that can benefit from the specializations of each other (e.g., divergence in access to resources, nutrient production capabilities and seasonal activity) (Walder et al. 2012). These findings, when combined with the observations of generalist mycorrhizae versus specialists, raise the question of what conditions favor each of these two mycorrhizal typesã and how do they affect the ecology of the forestã While much research is devoted to the direct mutual impact of trees and their inhabited mycorrhizal fungi, understanding the effect of mycorrhizal fungi symbionts on the interactions among trees and on tree-community is still limited.

This study has two aims: First, to map fungal-host associations in a mixed forest, while investigating the prevalence of generalism, namely those fungal species that inhabit various tree species and guilds of trees. Second, to study the potential implications of mycorrhizal fungi generalism versus specialism in that context. To do so, we identified the mycorrhizal fungal community in a mixed Mediterranean forest associated with four mature tree species which were previously shown to share C in a mesocosm experimental system (i.e. growing on the same soil; (Avital et al. 2022, Cahanovitc et al. 2022). Considering the high phylogenetic diversity of tree species in the mixed Mediterranean forest (including EM-hosts and AM-hosts), we hypothesized a high level of generalist mycorrhizal species overall (Hypothesis I). Next, we developed a stochastic simulation, modeling the competition dynamics between generalist and specialist mycorrhizal fungi in a mixed forest, incorporating tree-fungal interactions in the form of fungal-mediated resource sharing among trees. Using this simulation, we investigated the potential effect of resource sharing induced by generalist mycorrhizal fungi on forest diversity and structure. We hypothesized that when inter-tree species cooperation is beneficial, generalist mycorrhizal fungi that could mediate the resource sharing would grant heterogeneous neighboring trees an advantage, and thus promote forest diversity and spatial mixing (Hypothesis II). Conversely, a more diverse and spatially mixed forest would facilitate the proliferation of generalist mycorrhizal fungi.

## Materials and methods

### Study site

Mycorrhizal fungal community was studied in a mature forest research plot in Yishi Forest (part of Harel Forest), Judean Foothills, Israel (31º 43’
sN 34º 57’E, 320 m elevation) (Klein et al. 2013, Lapidot et al. 2019, Jakoby et al. 2020). The vegetation in this forest is dominated by the planted gymnosperm species *Pinus halepensis* and *Cupressus sempervirens* and the Mediterranean angiosperm woody species *Quercus calliprinos* and *Pistacia lentiscus*, accompanied by a variety of annual plants that thrive from winter to spring and several small woody species such as *Rhamnus lycioides, Phillyrea latifolia* and *Styrax officinalis* and *Ceratonia siliqua*, which belongs to the legume family. For brevity, we hereby refer to each of the four main species by their respective genus name, i.e., *Pinus, Cupressus, Quercus* and *Pistacia* (Due to the low sample numbers for *Ceratonia*, it was excluded from further analyses). In Yishi Forest, conifers were planted ~50 years ago, while broadleaf species populated the area before the plantation and soon after the conifer planting. The forest canopy is dominated by relatively tall conifers, especially pines, with lower stature *Quercus* and *Ceratonia*, and *Pistacia* forming a forest understory. *Pinus* and *Cupressus* are early succession species, featuring fast growth and massive seed production (Delipetrou et al. 2008, Sheffer 2012). In contrast, *Quercus, Ceratonia* and *Pistacia* exhibit late succession properties, such as a low growth rate, high shade tolerance and longer lifecycle (Ne’eman and Izhaki 1996, Bonet 2004, Maestre et al. 2004, Sheffer 2012). In terms of rooting depth, *Cupressus* and *Pistacia* dominate the shallow soil layer, *Quercus* and *Ceratonia* roots grow primarily in the rock layer, and *Pinus* roots inhabit both layers. The physical and chemical properties of the soil are detailed in Table **S1**. Using an eTrex30x GPS instrument (Garmin, Olathe, Kansas, USA), we mapped all the trees exceeding 1 m in height. Within the 1 hectare forest research plot, there were 242, 146, 109, 31, and 9 *Pistacia, Cupressus, Quercus, Pinus*, and *Ceratonia* trees, respectively. Our site offered a matrix design of tree and mycorrhizal functional types: EM-hosts included the coniferous *Pinus* and broadleaf *Quercus*, and AM-hosts included the coniferous *Cupressus* and broadleaf *Pistacia*. In that way, there was little room to conifer-specialist and broadleaf-specialist mycorrhizal species, which were otherwise identified as important at higher latitudes (van der Linde et al. 2018).

### Root tip collection and DNA extraction

Lateral fine roots from four representative trees per studied species were sampled during December 2017-February 2018 (i.e., in the wet season, when fungal activity is higher; 20 tree roots in total). Four different locations in the forest plot were chosen, with all 4 tree species within a distance of 10 m from each other (each plot was around 100 m^2^). The lateral fine roots were sampled at a distance of 1-4 m from the stems they belonged to, at the topsoil, at 10 cm soil depth. The selection of the right tree species was ensured by following the roots to the stems they emanated from and also by DNA sequence analysis of the sampled roots (detailed below). Six different terminal root branchlets were sampled per tree, covering all major azimuths around the stem. For each root branchlet, we recorded the distance from and orientation to the source tree. Within 24 hours, the sampled roots were stored in an ice box, followed by removing the attached soil particles under tap water, after which the washed fine roots were placed in Petri dishes with a thin layer of unchlorinated tap water and stored at 4 ^°^C fridge, and then placed over a 1 cm grid, to allow for the unbiased sampling of 30 root tips (from every root branchlets, total of 3600 root tips), regardless of their size, form, and color. Root tips were picked from each root sample using a dissecting scope and tweezers that were flame-sterilized between each picking. In addition, we collected 15 root tips originating from 3 greenhouse saplings of each tree species, to serve as a positive control for our root-tips collection from field-based samples. Community analysis of these control samples appear in Avital et al., (2022). Individual root-tips were placed in 96-well plates pre-filled with 18 µl of Sigma XNAT DNA isolation buffer (Sigma-Aldrich, Buchs, Switzerland) and then stored at 4 ^°^C until the DNA was extracted with an Extract-N-Amp Tissue PCR kit (SIGMA XNAT2-1KT) according to the manufacturer’s standard protocol, with the following modification: The 96-well plates were incubated for 10 minutes at room temperature, and then for 4 min at 95 ^°^C using a thermo-cycler, before being mixed with the neutralizing inhibitor solution (15 µl) and stored at 4 ^°^C. PCR products were run on agarose gel and led to more than one clear band per root-tip in all the tree species (Fig. **S1**). These results led us to modify the sequence analysis from individual Sanger sequencing to Illumina sequencing of bulk root tips (detailed below; 30 root-tips of every terminal root branchlets were bulked together).

### Molecular identification of tree species

From the bulk of the 30 root tips of every lateral root (the same root tips of root-associated fungal ASVs identification), we amplified the tree ITS-2 region and verified our assessment of tree species identify. PCR reactions were performed using 1.25 units of Dream Taq DNA polymerase (Thermo Scientific) in 25 µl reaction volumes with 1 µl DNA extract and 1 µl of each of the primers ITS2-S2L and ITS2-S3R (Table **S2**) (Yao et al. 2010). PCR reactions were performed as follows: 1 min at 94 ^°^C, followed by 35 cycles of 30 sec at 94 ^°^C, 30 sec at 56 ^°^C, and 1 min at 72 ^°^C, and then a final cycle of 8 min at 72 ^°^C. Six µl of the PCR products were run on 1.5% agarose gel; PCR reaction samples with one clear band were purified by Exo-1 and rSAP and Sanger sequenced at the Biological Services Department (Weizmann Institute of Science, Rehovot, Israel), uni-directionally with the primer ITS2-S3R. The ITS2 sequences were BLAST manually in the NCBI data set. Only samples with a clear tree species identity (>95% identity query cover) were further analyzed.

### Root tips’ morph type and fungal genetic identification

Individual root tips were photographed and genetically identified for their fungal identity and tree species identity. Each root sample was placed in a petri dish, and several root tips were picked using a dissecting scope and tweezers and pictured using binocular (SMZ800, Nikon, Tokyo, Japan) and digital sight (DS-Fi2, Nikon). The root tips were then placed in pre-filled 96-well plates with Sigma XNAT DNA isolation buffer, as described below. For fungal identification, we used the ITS region with the primers ITS1F (Gardes and Bruns 1993) and ITS4 (White et al. 1990). PCR reactions were performed as in Rog et al. (2020). In brief: PCR reactions were performed using 1.25 units Dream Taq DNA polymerase (Thermo Scientific) in 25 µl reaction volumes with 1 µl DNA extract, 0.2 mM dNTPs, and 0.5 µl primer. The PCR program was: 1 min at 94 ^°^C, followed by 35 cycles of 30 sec at 94 ^°^C, 30 sec at 51 ^°^C, and 1 min at 72 ^°^C, and then one final cycle of 8 min at 72 ^°^C. PCR reaction samples with one clear band were purified by Exo-1 and rSAP and Sanger sequenced uni-directionally with the primer ITS4, at the Biological Services Department (Weizmann Institute of Science, Rehovot, Israel). Electropherograms of raw sequences were checked manually and only high-quality portions of the sequence were assigned ASV names by manually editing ambiguous bases and queried in sequence databases at PlutoF (UNITE, BLASTN 2.4.0) (Altschul et al. 1997).

### Molecular identification of root-associated fungal ASVs

Root tip identification was based on ITS2, which is part of the DNA barcode for fungi, using Illumina sequencing. Primers for the amplicon library were designed according to Taylor et al. 2016, which are biased to EMF in the Basidiomycota and Ascomycota and less specific to AMF. A two-step library preparation protocol was performed according to Nejman et al. (2020), with several modifications: The PCR reactions were performed using KAPA HiFi HotStart ReadyMix DNA polymerase (Hoffmann-La Roch, Basel, Switzerland) in 50 µl reaction volumes with 5 µl DNA extract (bulk of 30 root-tips from every lateral root) and 1 µl of each of the primers 5.8S-Fun (Taylor et al. 2016) and RD2-ITS4Fun (Table **S2**). The reverse primer consisted of the ITS4-Fun primer (Taylor et al. 2016) with the linker adapter RD2. PCR reactions were performed as follows: 2 min at 98 °C, followed by 35 cycles of 10 sec at 98 °C, 15 sec at 55 °C and 35 sec at 72 °C, and then one final cycle of 5 min at 72 °C. Second PCR reactions were performed using the same DNA polymerase in 50 µl reaction volumes with 1/10 (5 µl) of the first PCR reaction and 1 µl of each of the primers P5-rd1-5.8S-Fun and RD2-Barcode (Table **S2**). The forward primer consisted of the adaptor p5, the linker RD1 and the primer 5.8S-Fun. The reverse primer consisted of the adapter RD2 and the individual barcode (8 nucleotides) for every sample. PCR reactions were performed similarly to the first PCR, albeit with only 6 cycles. PCR amplicons were cleaned using Qiaquick PCR purification kit (Qiagen, Hilden, Germany) and then quantified fluorescently with the Qubit dsDNA HS kit (Life Technologies Inc., Gaithersburg, MD, USA). Libraries were quality checked for concentration and amplicon size using the Agilent 2100 Bioanalyzer (Agilent Technologies, Santa Clara, CA, USA) and size selected with AMPure magnetic beads (Beckman Coulter Inc., Brea, CA, USA). All the samples were sequenced in one amplicon using Illumina MiSeq with 300 bp paired-end reads (PE300_V3) in the Grand Israel National Center for Personalized Medicine (Weizmann Institute of Science, Rehovot, Israel).

### Bioinformatic analysis

The raw sequences of 135 root samples (120 mature trees + 15 control saplings) were demultiplexed and the adapters were removed together with the barcodes. The sequences were analyzed using amplicon sequencing dada 2 package v. 1.7.9 in R (Callahan BJ et al., 2016). Only sequences whose length exceeded 50 bp with a mean number of expected errors below 2 (maxN = 0, maxEE = c(2,5), minLen = 50, truncQ = 2) were deemed sufficient for further analysis. Paired-end sequences were merged using MergePairs function (dada2). Finally, a dereplication procedure for each site was performed independently (sample level), applied using derepFastq function, and then all the files (135 files, corresponding to 135 root samples) were combined into one single Fasta file in order to obtain one ASV data file and compare the fungal composition of all the samples. We removed singletons (min unique size = 2) and chimera sequences using the ‘removeBimeraDenovo’ function, and we clustered sequences (id = 0.97) and performed taxonomic assignation (id = 0.98) with the public reference databases from UNITE (v.7.2). Non-fungal ASVs were removed. Data normalization was achieved by rarefying to the lowest number of sequences among the samples (Fig. **S2**). We set the percentile cut-off under which samples were discarded at 10%. We retained only ASVs with at least 10 reads in all the samples (Table **S3**). Finally, we termed fungal species that were found to colonize more than a single host tree species as ‘generalist’. Since we did not define these fungi at the genet level, we acknowledge that they might be strictly regarded as ‘potentially generalist’. For simplicity, we term them ‘generalist’ hereafter.

### Functional group classification

We acknowledge the existing complexity in assigning fungal species into functional groups such as saprophytic vs. mycorrhizal, and ectomycorrhizal vs. arbuscular mycorrhizal. In the literature, AMF belong to the subphylum Glomeromycotina, while EMF are either in the phylum Basidiomycota or Ascomycota. In this study, we used the most updated classification available on the FUNGuild database (Nguyen et al. 2016). We do not purport this classification to be 100% accurate, but rather use it as a reference for our empiric observations, whether they support this classification or not. Based on updated publications, several unique ASVs classifications were discussed in the discussion section.

### Statistical analysis

After verifying the tree species’ identities using ITS2 DNA markers, the constructed data set was unbalanced (with 47, 35, 13, 11 and only 4 root samples for *Pistacia, Cupressus, Quercus, Pinus* and *Ceratonia*, respectively). Due to the preferential root growth of *Ceratonia* in the rock layer, and its consequent low sample numbers, it was excluded from further analyses. We used a the bootstrap resampling procedure (Dixon 1993) to produce a balanced ASV table with the minimal number of samples per species (i.e., 11). We repeatedly (1,000 times) selected 11 random samples per species and calculated the basic statistics on each of the balanced ASV tables (within the loop) and on the averaged balanced ASV table. To illustrate the main axes discriminating between tree species, a permutational MANOVA (PERMANOVA) was performed based on a Bray– Curtis dissimilarity matrix, using the distance and adonis functions embedded in the Phyloseq package (version 1.28) (Fig. **S3**). This procedure was followed by non-metric multi-dimensional scaling (NMDS) using the bootstrap method onthe averaged balanced ASV table.

### β-diversity calculation

The ratio between plot-scale and host-scale mycorrhizal species diversity (*β* diversity) was calculated and partitioned, as described below. *β* diversity can reflect two different phenomena, namely, nestedness of species assemblages and spatial species turnover (Baselga 2010, Livne-Luzon et al. 2017). Nestedness of species assemblages occurs when the communities of sites/samples with a smaller number of species are subsets of the communities of richer sites/samples. This phenomenon can reflect a non-random process of species loss driven by factors that promote an orderly disaggregation of assemblages, e.g., competitive exclusion. However, it can also reflect a non-random process of species gain, driven by differential dispersal and the establishment of the species’ capacity to form the community. Spatial species turnover occurs when the community of each sample has an exclusive set of species. It can reflect the replacement of some species by others as a consequence of environmental sorting or spatial and historical constraints. To estimate these two β diversity components, we used the samples of each of the four tree species. Hence, our estimates refer to nestedness and turnover components of β diversity that reflect dissimilarity among samples within each tree species. We partitioned β diversity using the multiple-sample dissimilarity method, implanted in the “beta.multi” function of the “betapart” R package (index used: Sorensen). The obtained nestedness (βSNE) and turnover (βSIM) components of the β diversity were presented as proportions (Fig. **S4**). To test for significant differences between the β diversity components of two tree types (shallow vs. deep rooting types), we used a permutation test based on a script written in R (version 3.1). Specifically, samples were randomly shuffled between the four tree species. Next, the absolute differences (two-tailed test), in the proportion of nestedness, between the two tree types were recorded. P-values were estimated as the proportion of cases in which the absolute differences obtained during the permutations (1,000 runs) were equal to or greater than those found in the original data set.

### Stochastic simulation model

We developed a stochastic simulation (using Python 3.6), modeling the interactions among hosts (trees) and their inhabited symbionts (mycorrhizal fungi), based on (Hauert and Doebeli 2004, Lewin-Epstein and Hadany 2020). The simulations followed a population of trees growing in a 2D-lattice of size 36 × 36 (a forest), where each site is inhabited by one of two tree types, *A*or *B*. For simplicity, we assumed that each tree can be colonized by one of three classes/types of mycorrhizal fungi: mycorrhiza α, which can colonize only *A* trees (specialist), mycorrhiza *β*, which can colonize only *B* trees (specialist), and mycorrhiza *γ*, which can colonize both *A*and *B* trees (generalist). A tree can be colonized by one type of mycorrhizal fungi at the most. In each step in the simulation, three processes occur: (i) mycorrhizal-fungi (hyphae) expansion; (ii) resource sharing and fitness evaluation; and (iii) tree death. In this study, we assume that mycorrhizal fungi that forms hyphae that connects neighboring trees, can mediate resource sharing among the trees (Avital et al. 2022, Cahanovitc et al. 2022).

During mycorrhizal-fungi expansion, a fungi of type *i* (∈ {*α, β, γ*}) colonizing a tree has a probability of *T*_*i*_ to grow and expand to a randomly chosen adjacent tree that is either free of mycorrhizae or colonized by a different mycorrhizal fungi type. In the latter case, if the fungi succeeds in expanding to the new tree (according to *T*_*i*_), it will replace the resident mycorrhizal-fungi and colonize the new tree. After mycorrhizal-fungi expansion, resource sharing takes place. We assume that if two adjacent trees are colonized by the same mycorrhizal fungi type, then a network of this fungi that connects the roots of the two trees will form. We assume that neighboring trees that are connected to the same hyphae network share resources (i.e., cooperate). We assume that a cooperating tree pays a fitness cost on each simulation step, and that the resources a tree provides are divided equally among all recipient trees. Thus, the fitness of tree *i* is:

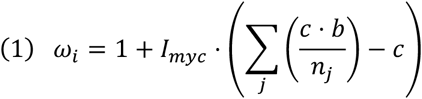

where *I*_*myc*_ equals 1 if tree *i* hosts mycorrhiza and is connected to at least one neighbor tree via the mycorrhiza, and 0 otherwise; *j* surveys the neighboring trees of *i* that host the same mycorrhiza type as *i*; *n*_*j*_ is the number of *j*’s neighbors to which j is connected via the same mycorrhiza type; *c* is the cost of cooperation; *b* is a benefit factor representing cooperation asymmetry, where resource sharing can benefit the recipient more than it costs the donor, for example, in situations where the recipient is less fit (e.g. stressed, young). The factor 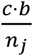 in the summation considers the benefit provided by tree *j* to tree *i*, which is the total benefits provided by tree *j* (*c*. *b*), divided by the number of neighbors among which these benefits are shared (*n*_*j*_).

We assume that all trees pay a fitness cost of *c* when cooperating, but the benefit of the recipient depends on the mycorrhizal species involved: *b*_α,*β*_ is the benefit to trees connected by mycorrhizas α or *β*, while for trees colonized by mycorrhiza *γ* the benefit depends also on the tree species: 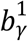is the benefit when *γ* connects two trees of the same species, and 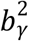is the benefit for trees of different species. In the main text we relate to 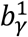 and 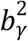 as the intra-tree-species benefit and to 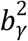as the inter-tree-species benefit. Finally, we included tree death: 5% of the trees in the forest die on each step, chosen with probabilities proportional to their fitness. If a dying tree is colonized by a mycorrhiza, the mycorrhiza dies with the tree. A seedling of one of the neighboring trees, chosen at random, will grow in the available site, starting free of any mycorrhizas. The fitness of all the trees resets to 1 at the beginning of each step.

## Results

### Shared and unique ASVs in the mixed forest

We analyzed 2,910 root tips of mature trees from four species: The planted conifers *Pinus halepensis* and *Cupressus sempervirens* and the local angiosperm woody species *Quercus calliprinos* and *Pistacia lentiscus*. Overall, we sampled 131 operational taxonomic units (ASVs) from 97 lateral roots belonging to the different hosts (42, 33, 12 and 10 root samples for *Pistacia, Cupressus, Quercus*, and *Pinus* trees, respectively) (Fig. **1a**). The abundance distribution of the various ASVs was log-normal, with a few common putative species (*Mycena maurella, Russula hortensis* and *Mycena capillaripes*) and many less abundant species (Fig. **1b**). Interestingly, although we detected a few saprotrophs and AMFs, the most abundant ASVs were EMFs (Fig. **1b**). The four most prevalent EMF ASVs were *Russula hortensis, Inocybe f. multifolia, Inocybe roseipes* and *Helvella lacunosa*. Our examination of the most abundant ASVs (i.e., those with a relative abundance larger than 10 in all the root samples, regardless of the host type) found two ASVs that were shared among all four host tree species (e.g., *Inocybe f. multifolia* and *Tricholoma terreum*, Fig. **1c**). Several ASVs were shared among three of the tree species (e.g., *Russula hortensis, Inocybe f. multifolia* and *Tricholoma terreum*) (Fig. **1a**), while others appeared only on the roots of a single host (e.g., *Mycena capillaripes* on *Pistacia*, and *Helvella lacunosa* on *Quercus*).

**Figure 1.**
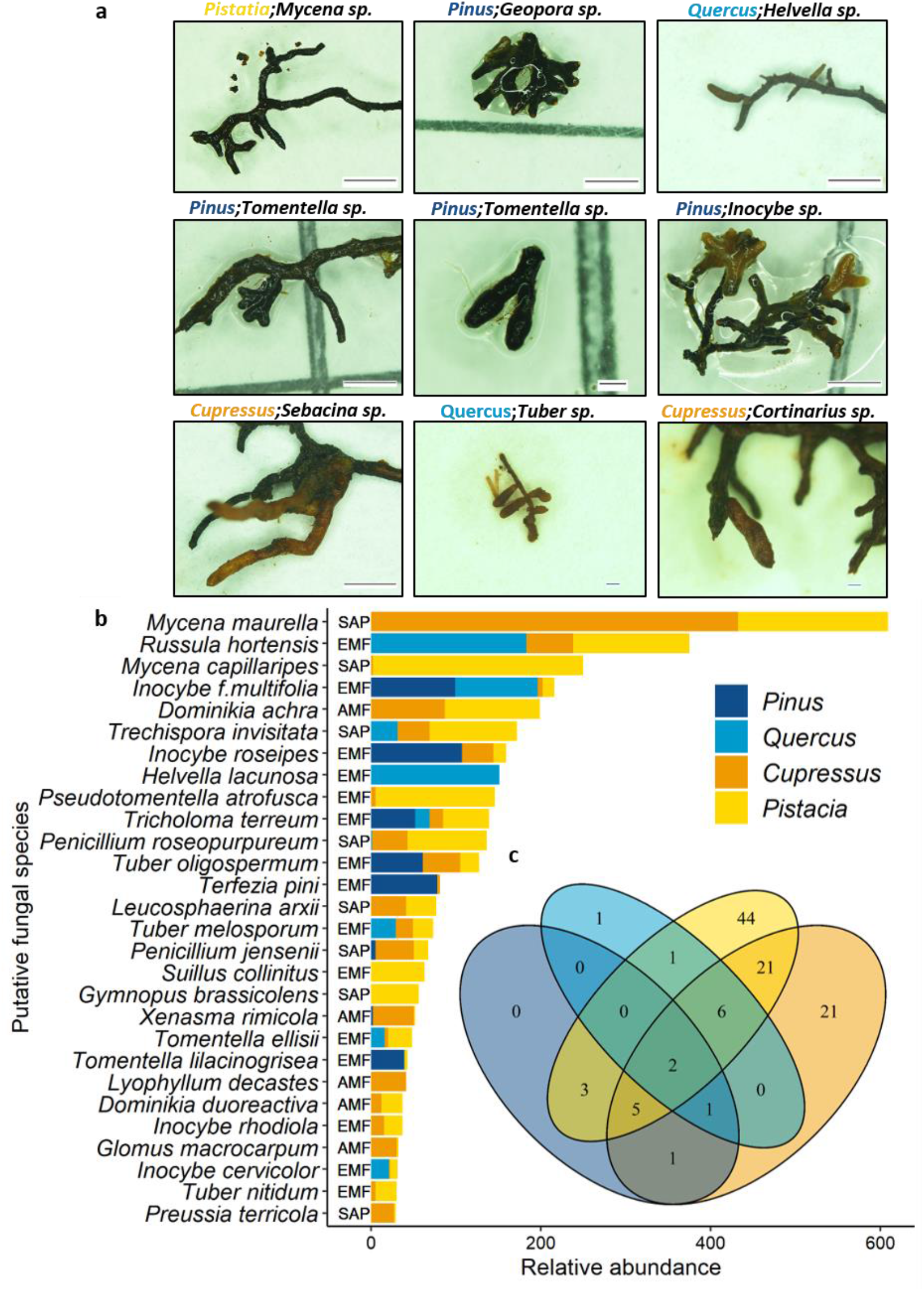
(**a**) Representative fungal and tree species’ root tip morphtype. The tree and fungal identification is based on the ITS2 and ITS4 sequence, respectively. Scale bars: 1 mm. (**b**). Stacked bar plot of the relative abundance of putative fungal species (higher than 20 reads after rarefaction). Colors represent host tree species; capital letters represent the association type according to FUNGuild (SAP-saprotrophic; EMF-ectomycorrhizal fungi; AMF-arbuscular mycorrhiza fungi). (**c**) Venn diagram displaying the number of unique and shared OUTs in root samples from four tree species.

### Mycorrhizal fungal community composition in the mixed forest

We used a bootstrap resampling procedure (see Methods) to produce a balanced ASV table of the mycorrhizal fungal community in the mixed forest. After the resampling procedure, the three most abundant ASVs were *Russula hortensis, Helvella lacunosa* and *Mycena maurella*, which were all located in a significantly separated dendrogram cluster chain of the mycorrhizal matrix (Fig. **2a**). Illustrating the main axes discriminating between tree species (permutational MANOVA (PERMANOVA) based on the Bray-Curtis dissimilarity matrix), the mycorrhizal community composition of the four tree species seems to split into three groups, where *Pistacia* and *Cupressus* closely relate while *Pinus* and *Quercus* stand apart (Fig. **2a**). For example, the major EMF genera *Tuber, Inocybe* and *Tomentella* colonized both *Pinus* and *Quercus* roots, while other species of these genera colonized either one of the tree species, namely *Pinus* or *Quercus*. Non-metric multi-dimensional scaling (NMDS) using a bootstrap method of the averaged balanced ASV table indicated that the mycorrhizal community of *Cupressus* and *Pistacia* differed from those of *Pinus* and *Quercus* (Fig. **2b**). The mycorrhizal communities of the classical EMF-host trees (*Pinus* and *Quercus*) and the classical AMF host trees (*Cupressus* and *Pistacia*) were significantly different, yet, unexpectedly, shared mycorrhizal species were found as well.

**Figure 2.**
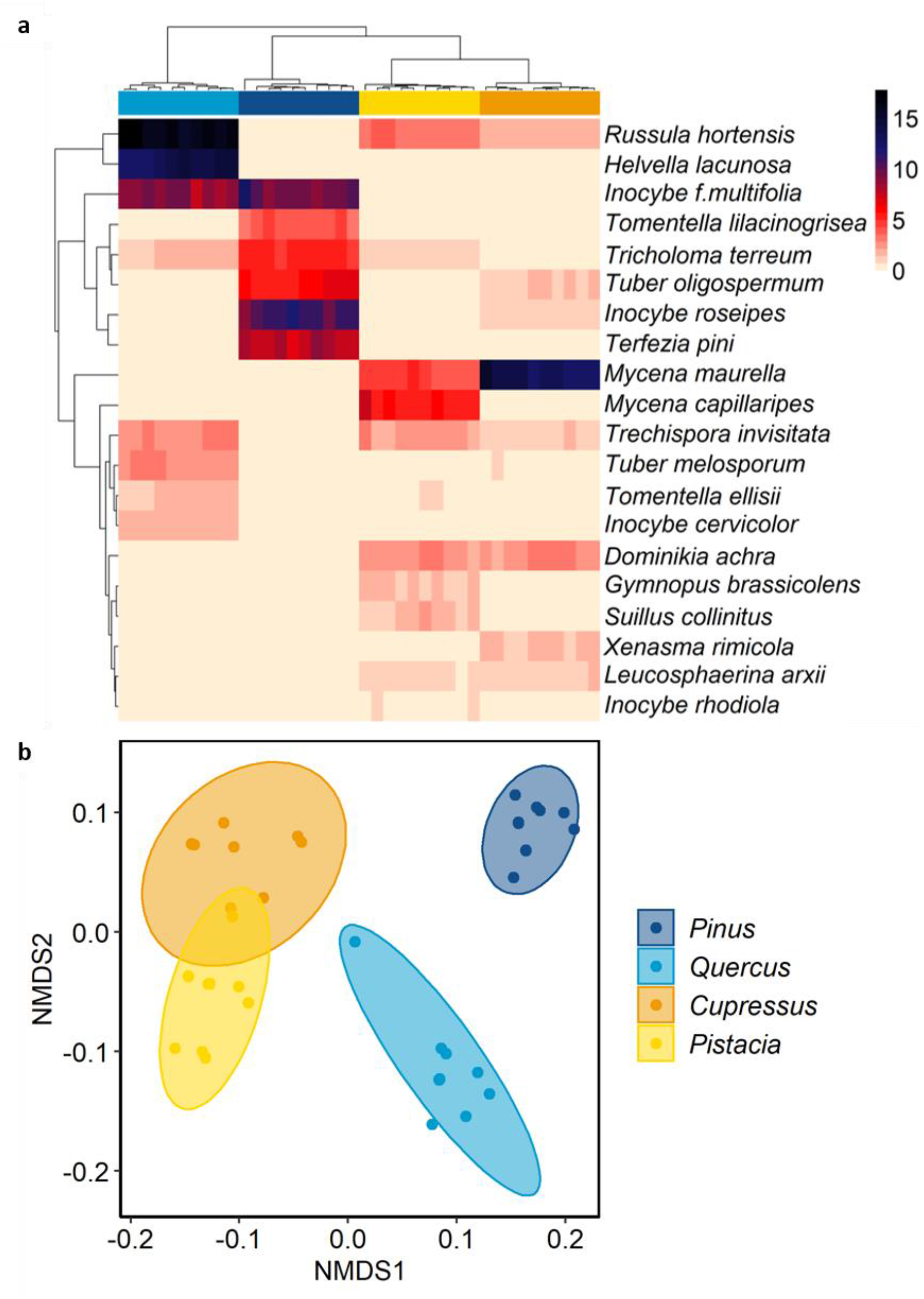
(**a**). Heat map of the community structure at the putative fungal species level (rows) by host tree species (columns) in terms of bootstrap average abundance (color coded). Dendrograms are constructed with Bray-Curtis distances. (**b**) The community composition of root-associated fungi, illustrated using non-metric multi-dimensional scaling (nMDS) of host tree species bootstrap averages. Circles represent 95% confidence intervals.

### Ectomycorrhizal fungi in the mixed forest connect to non-classical hosts

Beta diversity can reflect two different phenomena, nestedness of species assemblages and spatial species turnover. A permutation test applied to a resampled balanced data set (n=44) revealed significant differences (p<0.01) in the ASV nestedness component between the EMF-host and AMF-host tree guilds (Fig. **S4**). The classical EMF-host tree species exhibited a higher level of nestedness, while the AMF hosts displayed a higher rate of ASV turnover. Furthermore, we did not detect even one AMF species on the roots of the EMF-host trees (*Pinus* and *Quercus*) (Figs. **S3, S4**). In contrast, a number of EMF species were found to associate with known AMF-host trees (*Cupressus* and *Pistacia*) (Fig. **S5**). EMF ASVs such as *Russula hortensis, Inocybe f. multifolia* and *Inocybe rosepies* were found to be highly abundant and to be present in the two host guilds. Still, other EMF ASVs, such as *Helvella lacunosa, Terfezia pini* and *Tomentella lilacinogrisea*, were less abundant and were associated with a smaller number of hosts, mainly the classical EMF-host trees, with several others, like *Suillus collinitus, Inocybe rhodiola, Tomentella ellisi* and *Tuber nitidum*, present only on the roots of classical EMF-host trees. Overall, generalist EMF species were shared among the four tree species, with high abundance on EMF-host tree roots and low, but significant, abundance on AMF-host tree roots.

### Generalist-specialist competition model

We developed a computational model and constructed a stochastic simulation based on previous research (Hauert and Doebeli 2004, Lewin-Epstein and Hadany 2020) in order to study mycorrhiza-induced cooperation among trees (see Methods). Within this context, we focused on the dynamics that arise from the competition between specialist mycorrhizal fungi, that can inhabit and mediate resource sharing among a narrow set of trees, and generalists, that can inhabit and mediate resource sharing among a wide set of trees. We utilized the simulation to investigate the evolutionary dynamics between generalist and specialist mycorrhizal fungi and the potential impact on the forest diversity and spatial structure. We analyzed the benefits that mycorrhizal fungi can obtain by mediating resource sharing among its tree hosts, the conditions under which being a generalist (multi-host) fungi that can mediate resource sharing among different tree types is advantageous, and what may be the implications of proliferation of generalist fungi to the forest ecology.

The simulations modeled a 2D rectangular forest plot consisting of two tree species, A and B, where each tree can be colonized by one of three mycorrhizal types: mycorrhiza α which can colonize only A trees (specialist), mycorrhiza *β* which can colonize only B trees (specialist), and mycorrhiza *γ* which can colonize both A and B trees (generalist). In the context of our mixed forest, trees A and B can represent EMF and AMF tree hosts, respectively, while mycorrhiza α, *β, γ* can represent EMF colonizing only EMF tree hosts, AMF colonizing only AMF tree hosts, and EMF colonizing both EMF and AMF tree hosts, respectively. Following studies that demonstrated trees’ resource sharing mediated by mycorrhiza (Klein et al. 2016, Rog et al. 2020), and specifically in the system studied here (Avital et al. 2022, Cahanovitc et al. 2022), we assumed in our model that whenever two adjacent trees are colonized by the same mycorrhizal fungi type, a mycorrhizal network is formed, enabling resource sharing between the two trees. We considered the cost to the donor tree and the benefits to the recipients, mediated by the mycorrhizal fungi (distinguishing between intra- and inter-tree-types benefits), as well as the expansion rates of the different mycorrhizal fungi types – the rate at which the hyphae grows and expands from one colonized tree to a neighboring tree, and colonizes it too (Fig. **3a**).

**Figure 3.**
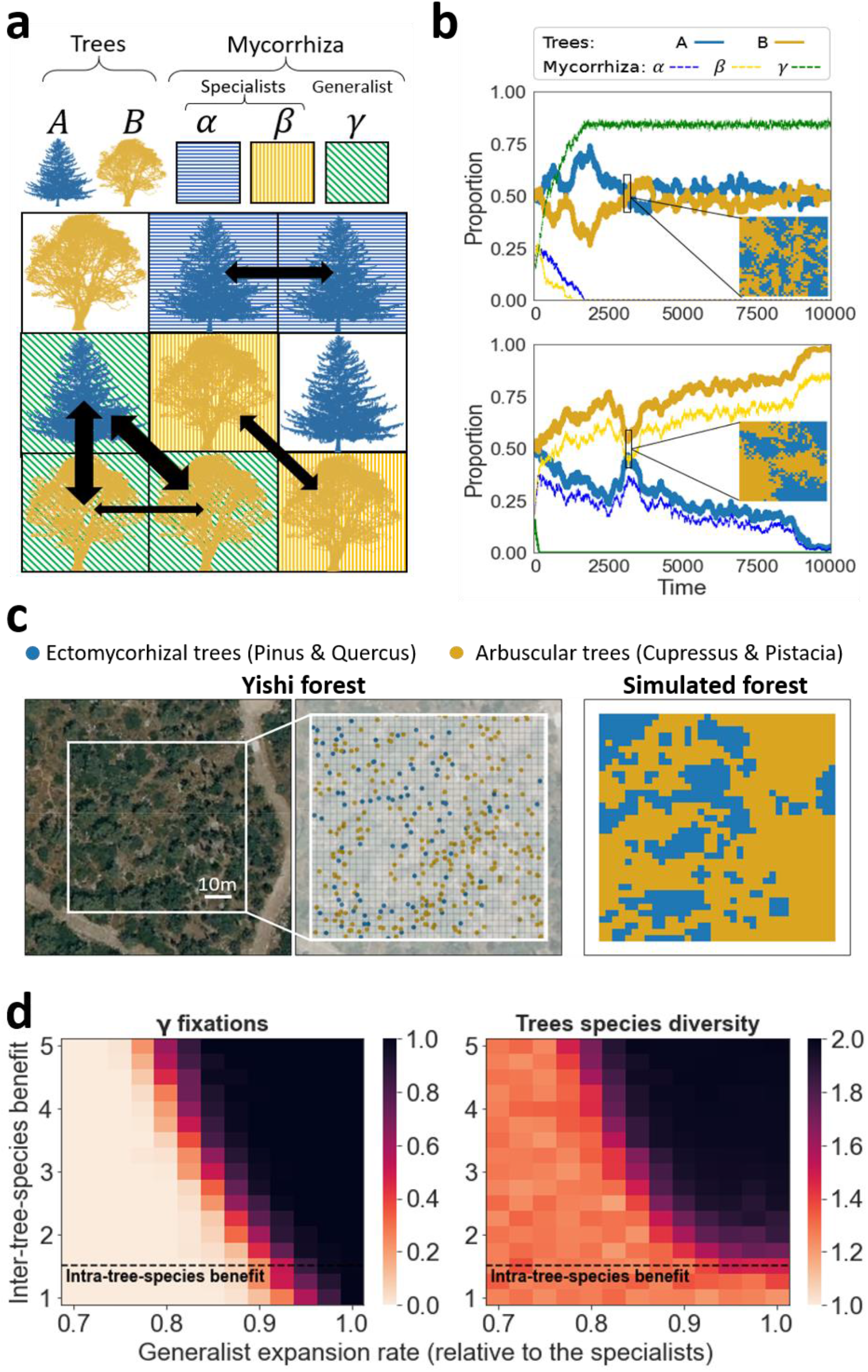
A generalist vs. specialist mycorrhiza competition model. (**a**) Illustration of the interactions within a simulated forest containing two tree species, A and B, and three mycorrhiza species: α, that can colonize only A trees (specialist); *β*, that can colonize only B trees (specialist); and *γ*, that can colonize both A and B trees (generalist). Arrows between trees represent resource sharing (see Methods). (**b**) Time series of two simulation runs where mycorrhiza *γ* either fixates (upper panel) or goes extinct (lower panel). Plotted are the changes in the proportions of the different mycorrhiza and tree species along time. The blue-yellow inset lattices present the structure of the forest at *t =* 3,150, with each pixel representing one tree. Although the proportion of both trees in both forests in *t =* 3,150 is close to 0.5, the spatial mixing level is significantly higher in the presence of mycorrhiza *γ*: ~0.31 (upper panel) and ~0.19 (lower panel). Inter tree species benefit was set to 3, the cost of cooperation to 0.2, the specialists’ (α, *β*) expansion rate to 0.1, and the generalist (*γ*) expansion rate to 0.09 (upper panel) and 0.08 (lower panel). The simulations were initialized with randomly located trees with equal proportions of both species, while half of the trees were colonized by a mycorrhiza, with equal proportions of all three species. (**c**) Generating a simulated forest based on a plot from Yishi forest. Trees in the plot were categorized as either EMF or AMF, and the plot was divided to sections with a 36 × 36 grid. Each tree in the simulated lattice was chosen to be A (EMF) or B (EMF) according to the type of the closest tree in the plot. (**d**) The expected proportion of *γ* (left panel) and the tree-species diversity (right panel; obtained by calculating 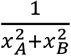, where *x*_*A*_and *x*_*B*_ are the proportions of trees *A*and *B*) after 10,000 time points, as function of the inter-species cooperation benefit (y-axis) and of the generalist (*γ*) expansion rate relative to that of the specialists (α, *β*; x-axis). The simulations were initialized with forest structure as presented in (c). Half of the trees were initially colonized by a mycorrhiza, with proportions as observed in the forest sampling: 10% α mycorrhiza (colonizing only EMF (A) trees); 50% *β* mycorrhiza (colonizing only AMF (B) trees); 40% *γ* mycorrhiza (colonizing both AMF and EMF (A and B) trees). Each data point represents the average of 100 simulations. The cost of cooperation was set to 0.2 and the specialists’ expansion rate to 0.1.

We found that the generalist mycorrhizal fungi (*γ*) can spread and take-over even when at an expansion disadvantage relative to the specialists (α, *β*; Fig. **3b**, upper panel) and even when the benefit of intra-species cooperation conferred by *γ* was lower than the benefit yielded by α or *β* (Fig. **S6**). Moreover, when *γ* proliferates, long-term species diversity among the trees can be maintained (Fig. **3b**, compare upper and lower panels). The competition dynamics between the different mycorrhizal species affects not only the diversity of tree species but also their spatial distribution in the forest. To quantify the forest structure, we defined the spatial mixing measure *ρ*(*t*) as the proportion of neighboring trees that are of a different type from the focal tree, averaged across all trees at time *t*. We found that even in cases where two forest plots had similar frequencies of trees A and B, the structure of a *γ*-dominated forest is much more mixed (Fig. **3b**, compare the insets).

We further validated these dynamics, initializing our simulation with tree distribution and mycorrhiza composition that was derived from the study plot in Yishi forest (Fig. **3c**), and investigating a wide range of parameters. We considered cases where the highest benefit of cooperation was between different species of trees (i.e., inter-species; for example, when trees can profit from each other’s specialization, such as access to resources and seasonal activity), and cases where the highest benefit was between trees of the same species (i.e., intra-species; for example, due to higher efficiency of the specialist mycorrhizal fungi). We found that the generalist mycorrhiza (*γ*) can spread in the population under a wide range of conditions (Fig. **3d**, left). In the cases where *γ* proliferates, if the benefit from inter-tree-species cooperation is higher than the benefit from intra-tree-species cooperation, then neighboring trees of different species are favored due to the *γ*-mediated beneficial resource sharing, and thus long-term species diversity among the trees is promoted (*γ*; Fig. **3d**). The advantage of *γ* results from two factors: (1) inter-species cooperation, and (2) host availability, as *γ* can colonize both A and B trees. Mycorrhiza *γ* can fixate even when the inter-species cooperation benefit is lower than that of the intra-species, due to the greater abundance of trees that can host *γ*. This occurs when the expansion rate of the generalist is close or equal to that of the specialists. Nevertheless, in this parameter range, long-term tree diversity is usually not maintained (Fig. **3d**, right, below the dashed line). When mycorrhiza *γ* goes extinct, a competition between mycorrhiza α and *β* and trees A and B takes place, eventually leading to the extinction of one of the tree-mycorrhiza couples, even in the absence of a selective advantage, due to drift (Fig. **3b**, bottom and Fig. **3d**), eventually yielding a homogenous forest.

Finally, we investigated the effect of the initial forest structure on the dynamics of both the trees and the mycorrhizal fungi. We ran simulations initialized with equal amounts of A and B trees, under four different configurations: 2, 4, and 6 homogenous patches of trees with equal proportions of the mycorrhiza species, initially colonizing half of the trees (Fig. **4a-c**), as well as an initialization that was derived from the study plot in Yishi forest (Fig. **4d**, Fig. **S7**; see derivation in Fig. **3**). We found that when the initial level of spatial mixing in the forest increases, mycorrhiza *γ* can evolve and fixate under a wider range of conditions, resulting in the maintenance of tree species diversity and higher levels of spatial mixing.

**Figure 4.**
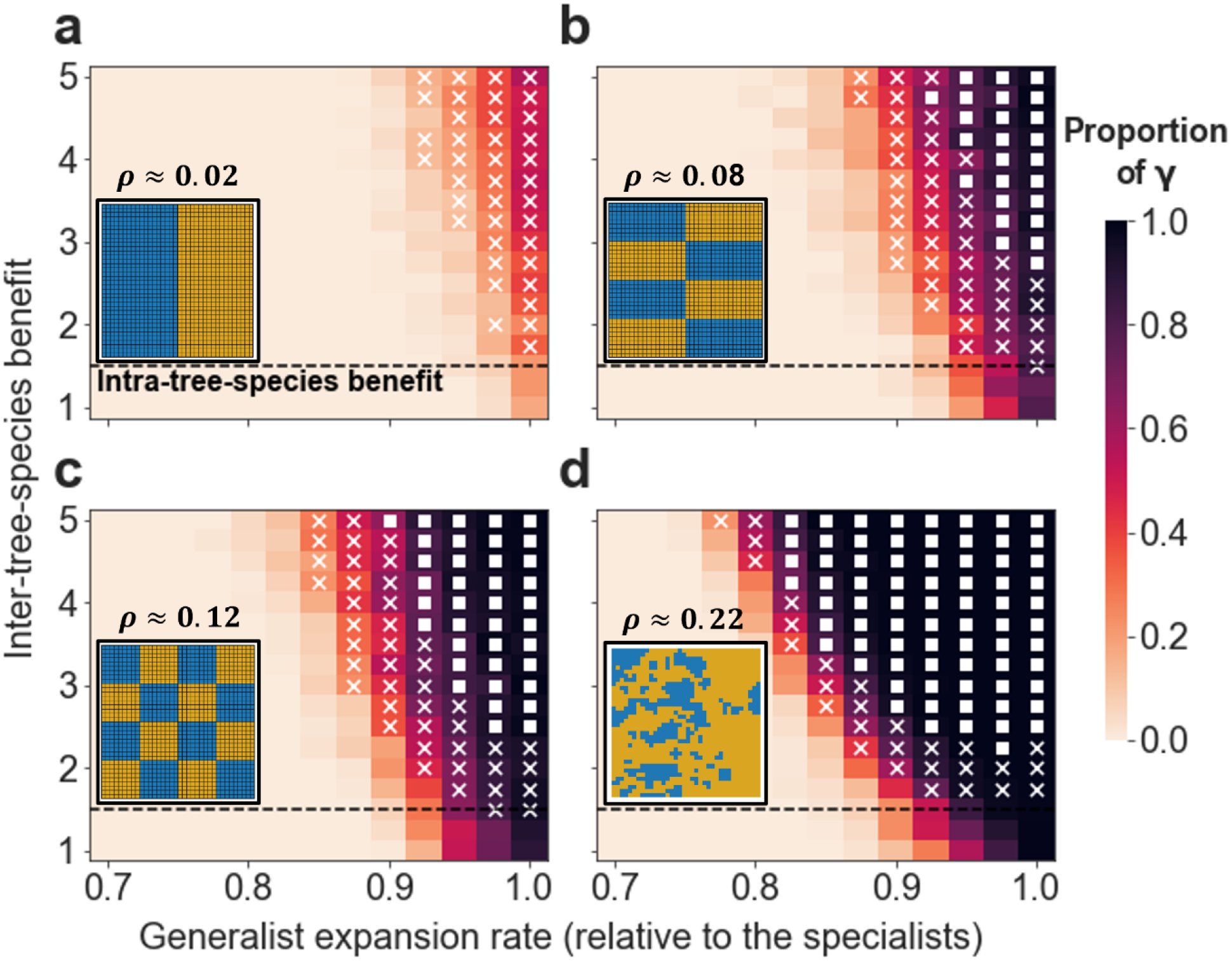
Higher initial trees spatial mixing allows the generalist mycorrhiza (*γ*) to evolve under a wider range of conditions. Plotted are the expected proportions of *γ* in the population after 10,000 simulation steps as a function of inter-species cooperation benefit (y-axis) and the generalist (*γ*) expansion rate relative to that of the specialists’ (α, *β*; x-axis). Insets indicate the initial locations of the A (blue) and B (yellow) trees in the simulations, while the specified *ρ* values represent the initial mixing level of the forest. In panel (**a**-**c**) the different mycorrhiza species occupy half of the trees with equal proportions. In panel (**d**) the forest is initialized based on a plot from Yishi forest as explained in Fig. 3. The x and square markings denote parameter ranges where the forest becomes spatially mixed after 10,000 simulation steps: instances where the trees in the forest have, on average, at least one (x marking; *ρ > 0*.*125*) or two (square marking; *ρ > 0*.*25*) neighbors that are of different species. The dashed lines indicate cases where the benefit of inter-species and intra-species cooperation are equal. Each data point in the plots represents the average of 100 simulations. In all the simulations, the cost of cooperation was set to 0.2 and the expansion rate of the specialist mycorrhiza species (α, *β*) to 0.1.

## Discussion

Our study of the root-associated fungal community in a diverse mixed forest shows multiple overlaps in the fungal community across different tree species, although none of the four tree species belongs to the same botanical family. These overlaps confirm Hypothesis I, i.e., a high level of generalism in our mycorrhizal community. Fungal communities of different tree hosts were highly diverse, with higher diversity in *Pinus* and *Quercus*, the known host trees of EMF. Several generalist EMF species were also found in association with *Pistacia* and *Cupressus*. Our observations are in line with reports of the highest fungal diversity in multi-tree species mixtures of EM-hosts and AM-hosts in a subtropical forest in China (Singavarapu et al. 2021). These results led us to a theoretic investigation of the potential impact of generalist mycorrhizal fungi to the ecology of the forest, focusing on mycorrhizal-induced resource sharing among trees. Using a computational model, we show that resource sharing between trees of different species, induced by generalist mycorrhiza, may play a crucial role in maintaining forest diversity and spatial mixing, investigating conditions that support Hypothesis II Hypothesis II.

### Mutualism host expansion in the mixed forest

The ecological and evolutionary mechanisms driving host specificity within a mixed forest are increasingly complex. Tree diversity and population dynamics are known to be affected by the resident mycorrhizal fungal community’s composition (Tedersoo et al. 2020), while the direct mechanisms and the mycorrhizal community characteristics and their effects on forest structure are less understood, although important. Recently, (Simard et al. 2021)showed that clearcutting decreased the cover of EM-hosts, while promoting AM-hosts in the post-harvest year. In this study, we examined the diversity of a mycorrhizal community in a forest plot containing a mixture of EMF- and AMF-host trees that live in close proximity and share many available resources (Rog et al. 2021b). We show that some “classical” AMF-host species, like *Cupressus* and *Pistacia*, associate also with several EMF species (e.g., *Inocybe* and *Tuber* species) (Figs. **1** and **2**). It is important to note that the “classical” separation between EMF and AMF hosts was well demonstrated in our study, since the high abundance of EMF fungi was associated with EMF hosts, and no AMF fungi were detected on non-AMF hosts (Fig. **2**). This exclusive directionality of host expansion, EMF on AMF-host, can be explained by the saprotrophic status of EMF versus the obligate biotrophic characteristic of AMF (Heklau et al. 2021). While our specific species of *Cupressus* and *Pistacia* were not yet confirmed as dual-mycorrhizal plants (i.e., AM arbuscules or coils and EM structures such as Hartig net are both observed on roots of the same plant), other species from the same genera are known for their ability to host both AM and EM fungi (Teste et al. 2020). Still, the fact that we detected EMF species on non-EMF hosts and C transfer among EMF host and AMF host of saplings from the same species (Avital et al. 2022), does not mean that these fungi function as mycorrhizal partners. In addition, decoupling between mycorrhizal abundance and function (i.e., fungi colonize few roots but transfer abundant resources versus fungi that colonize many roots but transfer limited resources through them) tangle the resource transfer via mycorrhizal networks (Bidartondo et al. 2001). The association between the EMF host and the AMF fungi requires further investigation, specifically in the characterization of the mutualistic association.

### Misdiagnosis of mycorrhizal guilds

The mixed Mediterranean forest in Israel is one of the unique places where AMF and EMF hosts have evolved to live in proximity for many years. At the single-plant level, several plants can function both as AMF and EMF hosts, either simultaneously within the same root system (Toju and Sato 2018), at different life stages or in different environments (Lapeyrie and Chilvers 1985, Chen et al. 2000). These ‘dual-mycorrhizal plant species’ have traditionally been considered uncommon and unusual (Lodge 2000). Some of these dual-mycorrhizal cases may be the result of the misdiagnosis of mycorrhizal status (Brundrett and Tedersoo 2019, Tedersoo et al. 2020). In their paper, Brundrett and Tedersoo (2019) mention numerous reasons for the misclassification of mycorrhizas, many of which were thoroughly addressed in this study. For example, we identified the host tree by DNA sequencing, not just by the morphological features of the root (Fig. **1**). We also Sanger sequenced some of the putative EMF root tips of AMF hosts and double-checked their identity (Fig. **1**). We used the most updated UNITE data (Abarenkov et al. 2010) while, further research will include specific data base for AMF (Delavaux et al. 2021). However, morphological identification such as AM arbuscules and EM Hartig net needs to be found on the same root to confirm the dual-colonization. The interaction between roots and fungi is not limited to symbiosis; hyphae can access additional roots without forming classical mycorrhizal morphology (Wang et al. 2021). This interaction can serve as a relay or hub to create well distributed mycorrhizal networks in the forest. *Pistacia* and *Cupressus* trees are well distributed around the forest (Fig. **3c**) having a localized advantage, possibly connecting distant trees throughout the mycorrhizal network. We confirmed the association type of mycorrhizal species using both traditional and advanced methods, though the ecophysiological activity of the EMF in their less classical hosts requires further research.

### Resource sharing supports the mutualistic niche theory

The mutualistic niche theory can describe the mycorrhizal mutualism contribution to the coexistence of plants within plant communities (Peay 2016). Most plants are able to uptake only a limited amount of inorganic nutrients from the soil, however, the interaction with mycorrhizal fungi can increase the diversity of organic compounds available for the plants and thus increase host performance and the host niche breadth. For example, increasing AMF richness was demonstrated to increase aboveground plant richness and productivity (Van Der Heijden et al. 1998). At the same time, the overlap in the mycorrhizal fungal community between co-occurring plant species can lead to functional convergence in nutrient uptake, rather than the expected differentiation. Mycorrhizal fungi have developed some local adaptation to host populations, which supports the hypothesis that local adaptation to host populations plays an essential role in host colonization evolution (Hoeksema and Thompson 2007). Equalizing mechanisms (Chesson 2000), such as mycorrhizal networks, are suggested to reduce competitive interactions between different hosts (Simard et al. 1997). While many host-mycorrhizal associations have evolved to be highly specific (Bruns et al. 2002), for mycorrhizal networks to emerge in a diverse forest system, the interaction among host trees and their mycorrhizal partners requires both the host and the fungi to expand their compatibility. From the host point of view: *Suillus* species, known to exhibit high host specificity, have been shown to associate with both their classical Pinaceae host and other, less traditional hosts, like *Quercus*, only in the presence of a co-planted Pinaceae host (Lofgren et al. 2018). In addition, dual-mycorrhizal plants can switch between EMF and AMF mycorrhizal mutualists under a wide range of biotic and abiotic conditions were characterized (Teste et al. 2020). Considering that EMF and AMF represent plant adaptation to slow and fast nutrient cycling rates, respectively(Averill et al. 2019), dual-mycorrhization might provide plasticity under fluctuating nutrient availability. From the fungal point of view: certain fungi exhibit a metabolic route change from saprotrophic to biotrophic metabolism. Some species of *Mycena* and *Gymnopus* are known as saprotrophic fungi that were recruited as mycorrhizal symbionts of certain mycoheterotrophic orchids(Selosse et al. 2010). *Mycena*, root-associated saprotrophic fungi, were found to occupy tree roots and to be involved in carbon and phosphorous transfer between fungi and their host in a manner similar to that of mycorrhizal species (Thoen et al. 2020, Harder et al. 2021). Here, the two *Mycena* species and the single *Gymnopus* species were identified in FUNGuild as saprotrophic (Fig. 1).

### Mycorrhizal-induced cooperation and forest ecology: theoretical model

The computational model we constructed incorporates variability in the ability of different mycorrhizal fungal species to associate with several tree species, as well as for the ability of mycorrhizal fungi to mediate resource sharing between the tree hosts (Avital et al. 2022, Cahanovitc et al. 2022). We found that generalist mycorrhizal fungi can evolve even when subjected to an expansion rate disadvantage, provided that inter-tree-species cooperation is beneficial enough (Fig. **3**). We further found that proliferation of multi-host mycorrhizal fungi that can induce inter-tree-species cooperation, combined with the greater benefit of inter-tree-species cooperation (compared to intra-tree-species) leading to a higher mixing-heterogeneity of trees’ spatial distribution (mixing level) in the forest. Generalist mycorrhiza can also proliferate in cases where the inter-species cooperation benefit is lower than that of the intra-species, due to higher availability of hosts, yet in these cases the diversity of the forest is not expected to be maintained.

Previous studies have investigated the benefits of interactions between trees and their mycorrhizal partners to the trees and mycorrhiza alike (Birch et al. 2020, Sapes et al. 2021), while others focused on forest structure and diversity, without taking mycorrhizas into account. In this study, we combine these approaches, by considering the ability of a mycorrhizal species to induce its host trees to share resources and assessing the implications of this mycorrhizal-induced tree cooperation to the ecology of the forest. This analysis indicates the important role mycorrhiza may play in maintaining a diverse, cooperative forest, which can increase forest resilience to future stresses (Lin 2011, Oliver et al. 2015, Levine et al. 2016, Obolski et al. 2017, Gillespie et al. 2020). In the current model, we made several simplifying assumptions. First, we assumed that mycorrhizae do not pass to a tree’s offspring through the seeds, nor stay in the ground when a tree dies. As has been previously shown (Lewin-Epstein et al. 2017, Lewin-Epstein and Hadany 2020), the ability of microbes to transmit not only between interactors but also from parent to offspring can further increase the benefit of cooperation-inducing microbes. If some mycorrhizal species indeed have the ability to disperse with the seeds, we would expect natural selection to favor an even greater effect of the mycorrhiza on its tree host, and specifically over its resource-sharing-between-trees behavior. In addition, in the current study, we did not consider a cheating behavior, nor the resistance of trees to mycorrhizal intervention in their resources. Nevertheless, these two aspects have been thoroughly investigated (Lewin-Epstein et al. 2017, Lewin-Epstein and Hadany 2020), demonstrating that microbe-induced cooperation can evolve even when facing these two phenomena.

### EM-host or AM-host, why not bothã The advantages of a generalist mycorrhiza

In our forest plot, spatial tree species distribution was close to random, allowing for the formation of common mycorrhizal networks among EM-hosts. Surprisingly, in our mixed Mediterranean forest, half of the sampled root-tips in the topsoil were colonized by EM fungi, while only ~30% of the trees are known as EM hosts (Fig. **3c**). However, 80% of the EM fungi (relative abundance) were colonizing both EM-hosts and AM-hosts (Fig. **1**). This relatively high abundance of EM fungi in the upper soil layer is striking given the contrasting root distribution with EM-hosts having deeper roots compared to the shallower AM-hosts (Rog et al 2021b). Both EM and AM mycorrhizal fungi increase host nutrient and water uptake, however the beneficial effect of simultaneous colonization by the two mycorrhizal types, to both the host and the fungi, is not well understood. Overall, there are more empirical evidence reporting the advantage or neutral effect on plant host responses of dual-colonization than a single type of interaction (Teste et al. 2020). The possible benefits of generalist fungi, connecting simultaneously to both types of hosts, can be related to resource partitioning among different types of hosts. For example, at the regional scale, water availability appeared to explain well the association of the known dual-mycorrhizal hosts, *Populus*, with EM fungi (Karst et al. 2021). At a smaller spatial scale, spatial partitioning of soil depths (Neville et al. 2002), can be a critical advantage in the mixed Mediterranean forest (Rog et al. 2021b). Specifically, our EM-hosts *Pinus* and *Quercus* are the deeper rooted species, offering a potential advantage for the more shallow rooted AM-hosts by connecting to the EM network.

Colonization of both types of trees can be advantageous also considering the tree species phenology, in turn affecting the capacity of the host to serve as a carbon source (Rog et al. 2021a). For example, *Pinus* and *Quercus* are more active in the wet than the dry season, while *Pistacia* is active more or less year-round, and hence connecting to both types could maximize carbon source along the year. Generalism among mycorrhizal species can be also understood as a risk management strategy, since no tree is immune to competition, disease, or death. While many of the above arguments are true also for any two hosts, being of similar or different type, a true generalist mycorrhiza (EM-AM-colonizer) can maximize these benefits. One also has to remember that these tree and fungal species have evolved in various settings; for example, a *Quercus*-*Pistacia* community is a more dominant vegetation type in the Levant than the mixed forest studied here. Our model simplified the complexity of the mycorrhiza community and specifically its species richness, focusing on modeling the dynamics between different types, or classes, of mycorrhizal fungi - specialist and generalist - in the context of their ability to mediate resource sharing among host trees. Focusing on these aspects, the model suggests conditions that could promote the proliferation of a generalist mycorrhiza type. Extension of the model to consider a rich mycorrhizal community may consider additional aspects such as multiple infections, separated niches, frequency dependent selection etc. In addition, the model results can integrate more empirical measurements of the mycorrhizal community’s seasonal variation and understanding of resource sharing in trees hosting multiple fungi colonization.

## Summary

Our findings demonstrate the wide occurrence of generalist mycorrhizal associations. The integration of host-microbe interactions, and specifically the microbe-induced cooperation, with specialist-generalist dynamics, highlights the potential role played by microorganisms in affecting the structure and composition of the host-community. This work suggests a new mechanism for promoting community diversity and mixing, which are prominent ecological attributes associated with the functioning and resilience of the community. Mycorrhizal fungi have been shown to mediate resource sharing among trees, and here we investigate the significance and implications of tree-tree interactions on forest structure and diversity. Further research is also required to quantify the empirical costs and benefits conferred to the cooperating trees and their recipients. It would also be intriguing to study empirically the differences in the extent of resource-sharing mediated by different mycorrhizal species, generalists or specialists alike.

## Acknowledgments

The authors thank Ofer Ovadia (BGU) and Hagai Shemesh (THC) for statistical analysis suggestions and general comments; Ravid Straussman, Lian Narunsky and Gili Rosenberg (WIS) for the NGS library preparation; Sydney Glassman (UCR) and Tom Bruns (UCB) for their help in primer design and mycorrhiza protocols; and Rotem Cahanovitc and Yaara Oppenhimer Shaanan (Weizmann Tree Lab) for lab assistance.

## Author contributions

IR performed all the empirical work under the supervision of TK. OLE performed all the modelling work under the supervision of LH. The bioinformatic and statistical analyses, functional group classification, and β-diversity calculation were performed by SLL and IR. All co-authors participated in the writing of this paper.

