## Supplementary information for "Prosperity of the commons: Generalist mycorrhizal species dominate a mixed forest and may promote forest diversity by mediating resource sharing among trees"

Supplementary Tables and Figures: Tables S1-S3; Figures S1-S6

**Table S1**. Soil's physical and chemical properties at different depths at Yishi Forest, Israel

| Depth (cm) | water in saturation (%) | pH | conductivity (dS/m) | Cl (mg $l^{-1}$) | Na (meq $l^{-1}$) | Ca (mg $l^{-1}$) | Mg (mg $l^{-1}$) | SAR (h) | N nitrate (mg $m^2$) | P Olsen (mg $m^2$) | $CaCl_2$ (mg $m^2$) | CaCO3 total (%) |
|---|---|---|---|---|---|---|---|---|---|---|---|---|
| 5 | 76 (1.5) | 7 (0) | 1.0 (0.04) | 34.03 (1.28) | 0.93 (0.033) | 75.83 (53.18) | 20.9 (2.35) | 0.39 (0.01) | 1.36 (0.35) | <3 | 45.1 (4.13) | 2.66 (0.33) |
| 15 | 70.3 (0.8) | 7.1 (0.1) | 0.75 (0.02) | 25.83 (1.22) | 0.86 (0.033) | 138 (7.51) | 6.93 (0.12) | 0.46 (0.028) | 2.03 (1.18) | <3 | 28.93 (0.73) | 3 (0.57) |
| 25 | 72 (2.5) | 7.1 (0.03) | 0.8 (0.1) | 29.9 (2.26) | 0.96 (0.066) | 173.4 (12.75) | 8.46 (1.18) | 18.27 (0.04) | 2.16 (0.91) | <3 | 31.96 (2.74) | 3.66 (0.66) |
| 35 | 73.6 (1.8) | 7 (0.1) | 1.09 (0.18) | 38.4 (8.65) | 1.1 (0.15) | 222.9 (49.88) | 14.23 (5.35) | 0.46 (0.04) | 2.13 (0.98) | <3 | 143.9 (4.2) | 6.66 (2.18) |

**Table S2**. Primer list used Mycorrhiza community identification and tree species verification

| Primer name | Sequence | Reference |
|---|---|---|
| ITS2-S2L | 5'-ATGCGATACTTGGTGTGAAT | Yao et al. 2010 |
| ITS2-S3R | 5'-GACGCTTCTCCAGACTACAAT | Yao et al. 2010 |
| 5.8S-Fun | 5'- AACTTTYRRCAAYGGATCWCT | Taylor et al. 2016 |
| RD2-ITS4Fun | 5'AGACGTGTGCTCTTCCGATCT-AGCCTCCGCTTATTGATATGCTTAART | Taylor et al. 2016 |
| P5-rd1-5.8S-Fun | 5'- AATGATACGGCGACCACCGAGATCT-ACACTCTTTCCCTACACGACGCTCTTCCGATCT-AACTTTYRRCAAYGGATCWCT | |
| RD2-Barcode | 5' AGACGTGTGCTCTTCCGATCT-BARCODE | |

**Table S3**. Number of lateral roots, number of root tips, success of root tree identification and number of identified ASV's per tree. Three saplings from the different tree species were used as control for the EM and AM hosts.

| Sample | Mature/Saplings | Tree Sp. prediction | Tree name | Lateral root number | Number of root tips | Tree Sp. DNA identification | Number of Identified ASV's |
|---|---|---|---|---|---|---|---|
| X1 | mature | *Cupressus* | 1 | 1 | 30 | *Quercus* | 1 |
| X2 | mature | *Cupressus* | 1 | 2 | 30 | *Pistacia* | 3 |
| X3 | mature | *Cupressus* | 1 | 3 | 30 | *Pistacia* | 2 |
| X4 | mature | *Cupressus* | 1 | 4 | 30 | *Pistacia* | 4 |
| X5 | mature | *Cupressus* | 1 | 5 | 30 | *Pistacia* | 2 |
| X6 | mature | *Cupressus* | 1 | 6 | 30 | *Cupressus* | 4 |
| X7 | mature | *Cupressus* | 4 | 1 | 30 | *other* | NA |
| X8 | mature | *Cupressus* | 4 | 2 | 30 | *Quercus* | 2 |
| X9 | mature | *Cupressus* | 4 | 3 | 30 | *Pistacia* | 1 |
| X10 | mature | *Cupressus* | 4 | 4 | 30 | *Quercus* | 2 |
| X11 | mature | *Cupressus* | 4 | 5 | 30 | *Quercus* | 2 |
| X12 | mature | *Cupressus* | 4 | 6 | 30 | *Pistacia* | 7 |
| X13 | mature | *Cupressus* | Alp | 1 | 30 | *Quercus* | 1 |
| X14 | mature | *Cupressus* | Alp | 2 | 30 | *Pistacia* | 1 |
| X15 | mature | *Cupressus* | Alp | 3 | 30 | *Pistacia* | 0 |
| X16 | mature | *Cupressus* | Alp | 4 | 30 | *Pistacia* | 0 |
| X17 | mature | *Cupressus* | Alp | 5 | 30 | *Cupressus* | 3 |
| X18 | mature | *Cupressus* | Alp | 6 | 30 | *Pistacia* | 0 |
| X19 | mature | *Cupressus* | Bet | 1 | 30 | *Quercus* | 2 |
| X20 | mature | *Cupressus* | Bet | 2 | 30 | *Quercus* | 2 |
| X21 | mature | *Cupressus* | Bet | 3 | 30 | *Quercus* | 3 |
| X22 | mature | *Cupressus* | Bet | 4 | 30 | *Quercus* | 2 |
| X23 | mature | *Cupressus* | Bet | 5 | 30 | *Quercus* | 2 |
| X24 | mature | *Cupressus* | Bet | 6 | 30 | *Quercus* | 0 |
| X25 | Saplings | *Cupressus* | G | 1 | 15 | *Cupressus* | 2 |
| X26 | Saplings | *Cupressus* | G | 2 | 15 | *Cupressus* | 3 |
| X27 | Saplings | *Cupressus* | G | 3 | 15 | *Cupressus* | 1 |

| | | | | | | | |
|---|---|---|---|---|---|---|---|
| X28 | mature | *Ceratonia* | 3 | 1 | 30 | *Cupressus* | 3 |
| X29 | mature | *Ceratonia* | 3 | 2 | 30 | *Pistacia* | 3 |
| X30 | mature | *Ceratonia* | 3 | 3 | 30 | *Cupressus* | 3 |
| X31 | mature | *Ceratonia* | 3 | 4 | 30 | *Cupressus* | 5 |
| X32 | mature | *Ceratonia* | 3 | 5 | 30 | *Cupressus* | 0 |
| X33 | mature | *Ceratonia* | 3 | 6 | 30 | *Cupressus* | 5 |
| X34 | mature | *Ceratonia* | A | 1 | 30 | *Cupressus* | 1 |
| X35 | mature | *Ceratonia* | A | 2 | 30 | *Cupressus* | 5 |
| X36 | mature | *Ceratonia* | A | 3 | 30 | *Cupressus* | 3 |
| X37 | mature | *Ceratonia* | A | 4 | 30 | *Cupressus* | 2 |
| X38 | mature | *Ceratonia* | A | 5 | 30 | *Cupressus* | 1 |
| X39 | mature | *Ceratonia* | A | 6 | 30 | *Cupressus* | 2 |
| X40 | mature | *Ceratonia* | B | 1 | 30 | *Ceratonia* | 9 |
| X41 | mature | *Ceratonia* | B | 2 | 30 | *Pistacia* | 5 |
| X42 | mature | *Ceratonia* | B | 3 | 30 | *Pistacia* | 1 |
| X43 | mature | *Ceratonia* | B | 4 | 30 | *Ceratonia* | 2 |
| X44 | mature | *Ceratonia* | B | 5 | 30 | *Ceratonia* | 4 |
| X45 | mature | *Ceratonia* | B | 6 | 30 | *Ceratonia* | 5 |
| X46 | Saplings | *Ceratonia* | G | 1 | 15 | *Ceratonia* | 5 |
| X47 | Saplings | *Ceratonia* | G | 2 | 15 | *Ceratonia* | 3 |
| X48 | Saplings | *Ceratonia* | G | 3 | 15 | *Ceratonia* | 3 |
| X49 | mature | *Pinus* | 1 | 1 | 30 | *Pistacia* | 4 |
| X50 | mature | *Pinus* | 1 | 2 | 30 | *Pinus* | 1 |
| X51 | mature | *Pinus* | 1 | 3 | 30 | *noSeq* | NA |
| X52 | mature | *Pinus* | 1 | 4 | 30 | *Pinus* | 2 |
| X53 | mature | *Pinus* | 1 | 5 | 30 | *Pistacia* | 3 |
| X54 | mature | *Pinus* | 1 | 6 | 30 | *Pistacia* | 2 |
| X55 | mature | *Pinus* | Alp | 1 | 30 | *Pinus* | 2 |
| X56 | mature | *Pinus* | Alp | 2 | 30 | *Cupressus* | 5 |
| X57 | mature | *Pinus* | Alp | 3 | 30 | *Pistacia* | 3 |

| | | | | | | | |
|---|---|---|---|---|---|---|---|
| X58 | mature | *Pinus* | Alp | 4 | 30 | *Cupressus* | 5 |
| X59 | mature | *Pinus* | Alp | 5 | 30 | *Pistacia* | 6 |
| X60 | mature | *Pinus* | Alp | 6 | 30 | *Pinus* | 3 |
| X61 | mature | *Pinus* | Bet | 1 | 30 | *Pinus* | 3 |
| X62 | mature | *Pinus* | Bet | 2 | 30 | *Pinus* | 0 |
| X63 | mature | *Pinus* | Bet | 3 | 30 | *Pinus* | 0 |
| X64 | mature | *Pinus* | Bet | 4 | 30 | *Pinus* | 3 |
| X65 | mature | *Pinus* | Bet | 5 | 30 | *Pinus* | 3 |
| X66 | mature | *Pinus* | Bet | 6 | 30 | *Pinus* | 1 |
| X67 | mature | *Pinus* | E | 1 | 30 | *Pinus* | 1 |
| X68 | mature | *Pinus* | E | 2 | 30 | *Pistacia* | 4 |
| X69 | mature | *Pinus* | E | 3 | 30 | *noSeq* | NA |
| X70 | mature | *Pinus* | E | 4 | 30 | *Pistacia* | 2 |
| X71 | mature | *Pinus* | E | 5 | 30 | *Pistacia* | 4 |
| X72 | mature | *Pinus* | E | 6 | 30 | *Pistacia* | 2 |
| X73 | Saplings | *Pinus* | G | 1 | 15 | *Pinus* | 3 |
| X74 | Saplings | *Pinus* | G | 2 | 15 | *Pinus* | 1 |
| X75 | Saplings | *Pinus* | G | 3 | 15 | *Pinus* | 3 |
| X76 | mature | *Pistacia* | 1 | 1 | 30 | *Cupressus* | 4 |
| X77 | mature | *Pistacia* | 1 | 2 | 30 | *Pistacia* | 5 |
| X78 | mature | *Pistacia* | 1 | 3 | 30 | *noSeq* | NA |
| X79 | mature | *Pistacia* | 1 | 4 | 30 | *Pistacia* | 6 |
| X80 | mature | *Pistacia* | 1 | 5 | 30 | *Pistacia* | 3 |
| X81 | mature | *Pistacia* | 1 | 6 | 30 | *noSeq* | NA |
| X82 | mature | *Pistacia* | Alp | 1 | 30 | *Pistacia* | 2 |
| X83 | mature | *Pistacia* | Alp | 2 | 30 | *Pistacia* | 2 |
| X84 | mature | *Pistacia* | Alp | 3 | 30 | *noSeq* | NA |
| X85 | mature | *Pistacia* | Alp | 4 | 30 | *Pistacia* | 4 |
| X86 | mature | *Pistacia* | Alp | 5 | 30 | *Pistacia* | 2 |
| X87 | mature | *Pistacia* | Alp | 6 | 30 | *Pistacia* | 3 |

| | | | | | | | |
|---|---|---|---|---|---|---|---|
| X88 | Saplings | *Pistacia* | G | 1 | 15 | *Pistacia* | 1 |
| X89 | Saplings | *Pistacia* | G | 2 | 15 | *Pistacia* | 2 |
| X90 | Saplings | *Pistacia* | G | 3 | 15 | *Pistacia* | 2 |
| X91 | mature | *Pistacia* | Gam | 1 | 30 | *Pistacia* | 5 |
| X92 | mature | *Pistacia* | Gam | 2 | 30 | *Pistacia* | 5 |
| X93 | mature | *Pistacia* | Gam | 3 | 30 | *Pistacia* | 0 |
| X94 | mature | *Pistacia* | Gam | 4 | 30 | *Pistacia* | 6 |
| X95 | mature | *Pistacia* | Gam | 5 | 30 | *Pistacia* | 6 |
| X96 | mature | *Pistacia* | Gam | 6 | 30 | *Pistacia* | 6 |
| X97 | mature | *Pistacia* | X | 1 | 30 | *Pistacia* | 3 |
| X98 | mature | *Pistacia* | X | 2 | 30 | *Pistacia* | 10 |
| X99 | mature | *Pistacia* | X | 3 | 30 | *Pistacia* | 8 |
| X100 | mature | *Pistacia* | X | 4 | 30 | *Pistacia* | 5 |
| X101 | mature | *Pistacia* | X | 5 | 30 | *Pistacia* | 5 |
| X102 | mature | *Pistacia* | X | 6 | 30 | *Pistacia* | 1 |
| X103 | mature | *Ceratonia* | 1 | 1 | 30 | *Quercus* | 0 |
| X104 | mature | *Ceratonia* | 1 | 2 | 30 | *Quercus* | 1 |
| X105 | mature | *Ceratonia* | 1 | 3 | 30 | *Pistacia* | 4 |
| X106 | mature | *Ceratonia* | 1 | 4 | 30 | *Pistacia* | 1 |
| X107 | mature | *Ceratonia* | 1 | 5 | 30 | *Pistacia* | 0 |
| X108 | mature | *Ceratonia* | 1 | 6 | 30 | *Pistacia* | 0 |
| X109 | mature | *Quercus* | 2 | 1 | 30 | *Cupressus* | 1 |
| X110 | mature | *Quercus* | 2 | 2 | 30 | *Cupressus* | 3 |
| X111 | mature | *Quercus* | 2 | 3 | 30 | *Cupressus* | 5 |
| X112 | mature | *Quercus* | 2 | 4 | 30 | *noSeq* | NA |
| X113 | mature | *Quercus* | 2 | 5 | 30 | *Cupressus* | 7 |
| X114 | mature | *Quercus* | 2 | 6 | 30 | *noSeq* | NA |
| X115 | mature | *Quercus* | Bet | 1 | 30 | *Cupressus* | 2 |
| X116 | mature | *Quercus* | Bet | 2 | 30 | *Cupressus* | 1 |
| X117 | mature | *Quercus* | Bet | 3 | 30 | *Pistacia* | 1 |

| | | | | | | | |
|---|---|---|---|---|---|---|---|
| X118 | mature | *Quercus* | Bet | 4 | 30 | *noSeq* | NA |
| X119 | mature | *Quercus* | Bet | 5 | 30 | *other* | NA |
| X120 | mature | *Quercus* | Bet | 6 | 30 | *Cupressus* | 1 |
| X121 | mature | *Quercus* | C | 1 | 30 | *Cupressus* | 0 |
| X122 | mature | *Quercus* | C | 2 | 30 | *Cupressus* | 0 |
| X123 | mature | *Quercus* | C | 3 | 30 | *Cupressus* | 1 |
| X124 | mature | *Quercus* | C | 4 | 30 | *Cupressus* | 2 |
| X125 | mature | *Quercus* | C | 5 | 30 | *Cupressus* | 2 |
| X126 | mature | *Quercus* | C | 6 | 30 | *Cupressus* | 3 |
| X127 | Saplings | *Quercus* | G | 1 | 15 | *Quercus* | 2 |
| X128 | Saplings | *Quercus* | G | 2 | 15 | *Quercus* | 1 |
| X129 | Saplings | *Quercus* | G | 3 | 15 | *Quercus* | 1 |
| X130 | mature | *Quercus* | Gam | 1 | 30 | *Cupressus* | 3 |
| X131 | mature | *Quercus* | Gam | 2 | 30 | *Cupressus* | 4 |
| X132 | mature | *Quercus* | Gam | 3 | 30 | *Cupressus* | 4 |
| X133 | mature | *Quercus* | Gam | 4 | 30 | *Cupressus* | 3 |
| X134 | mature | *Quercus* | Gam | 5 | 30 | *Cupressus* | 0 |
| X135 | mature | *Quercus* | Gam | 6 | 30 | *Cupressus* | 1 |

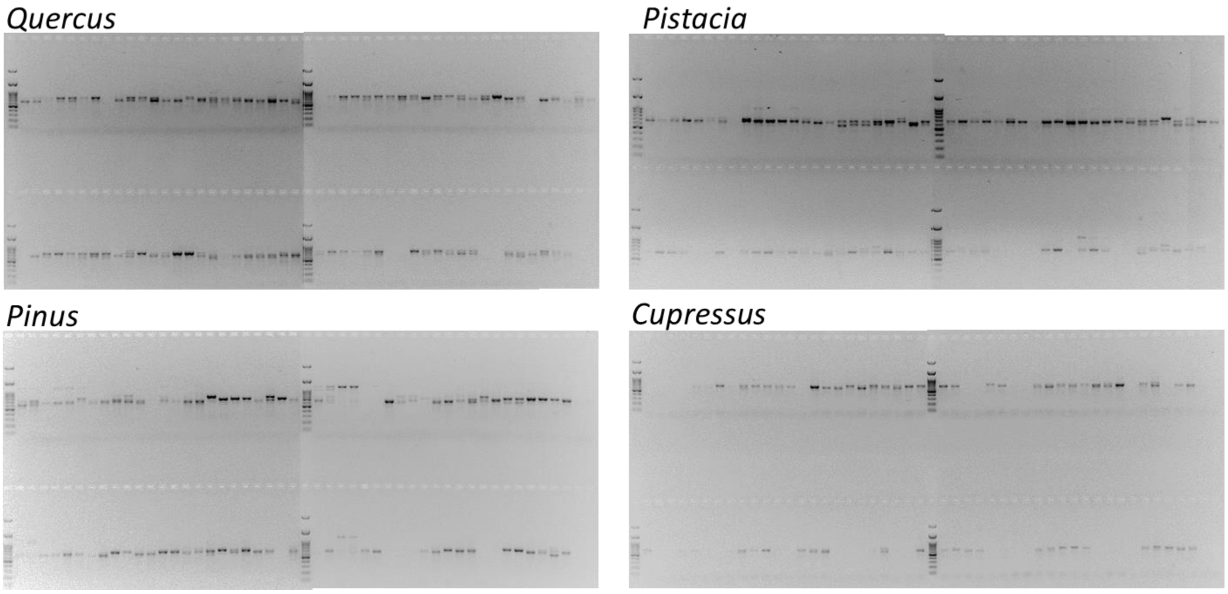


**Figure S1. PCR products from root tip of each tree species.** Agarose gel electrophoresis of PCR products generated using primers ITS1 and ITS4, from 30 root tips of each lateral root, 6 lateral root per tree.

a

b

c

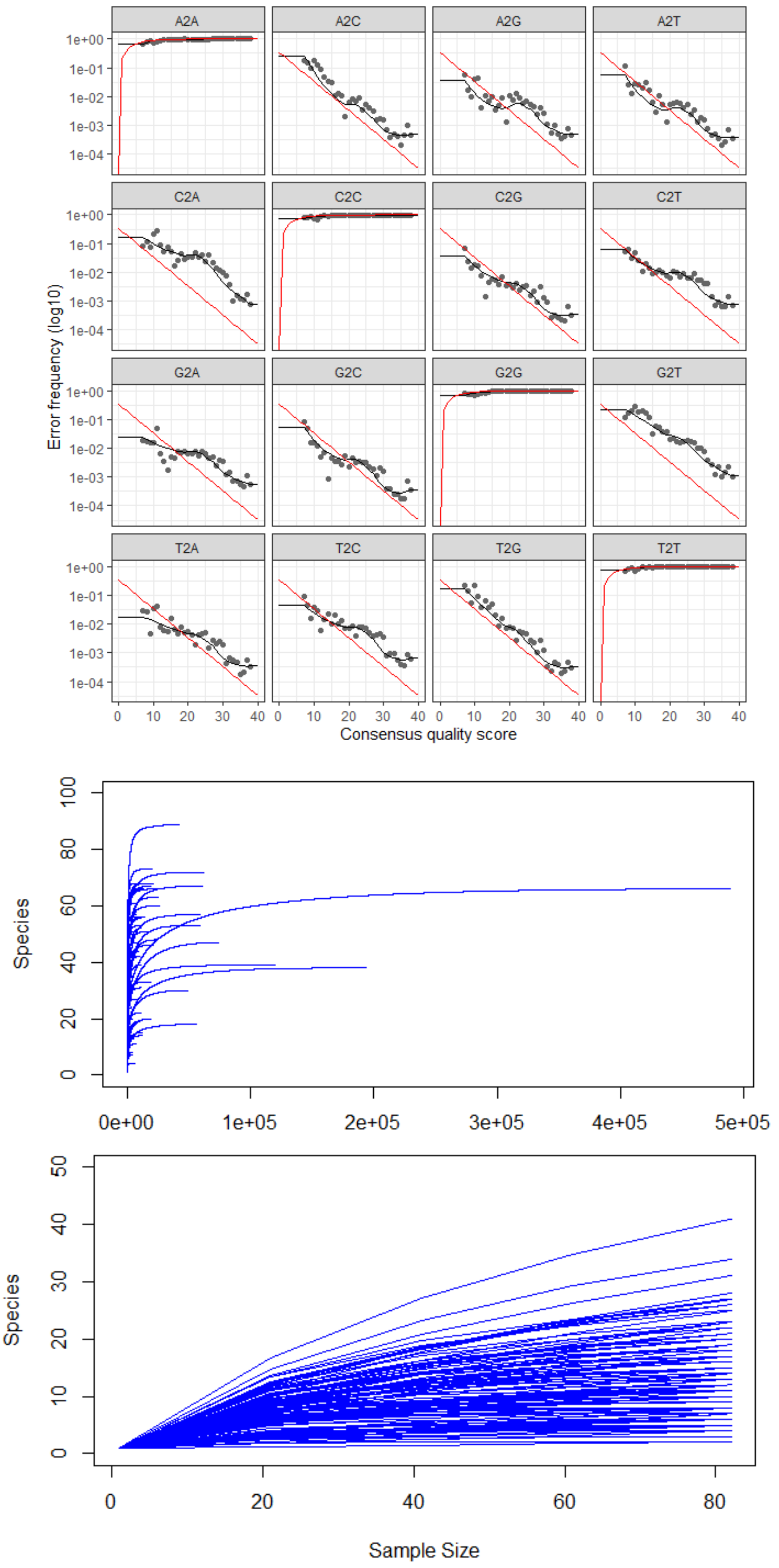


**Figure S2.** (a). Error frequency (b) before and (c) after rarefication.

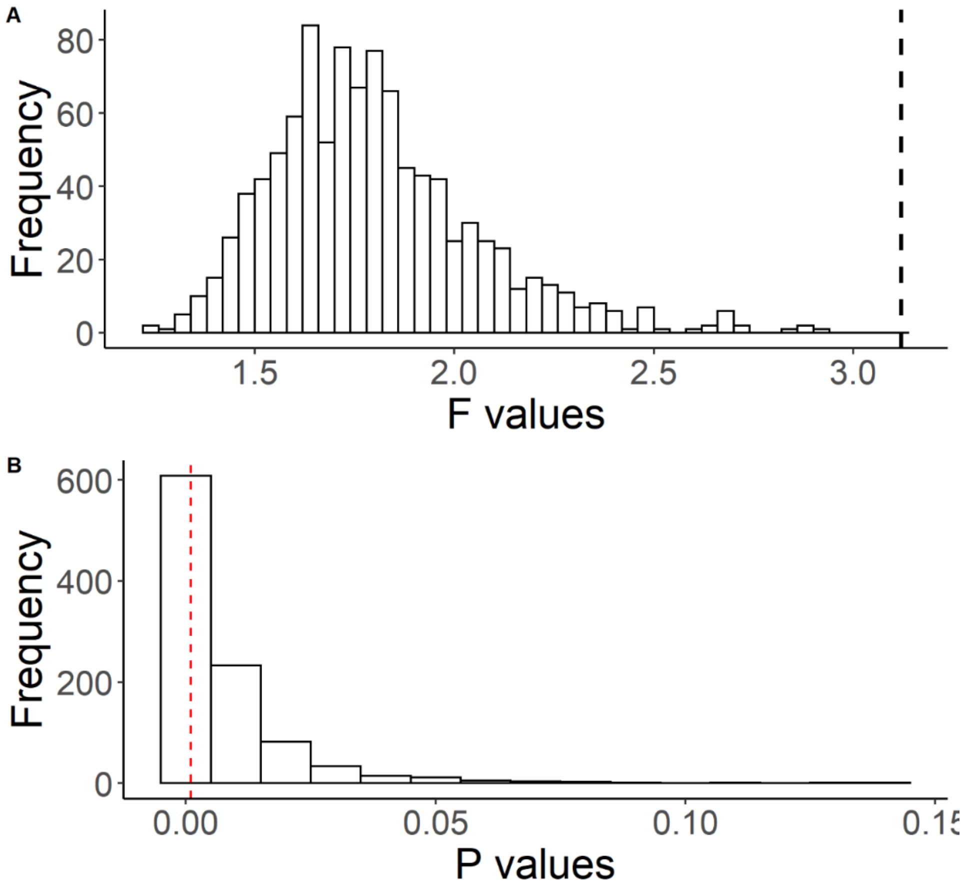


**Figure S3**. PERMANOVA test results: Histograms of (a) the F-statistics and (b) p-values of 1,000 tests that examined the differences in the OTU community of mature forest trees from a mixed forest plot using a resampled balanced data set (n=44). Dashed black lines denote the F and p values calculated based on the original unbalanced data set; the grey line denotes the p-value calculated based on the averaged OTU table of all the 1,000 balanced resamples.

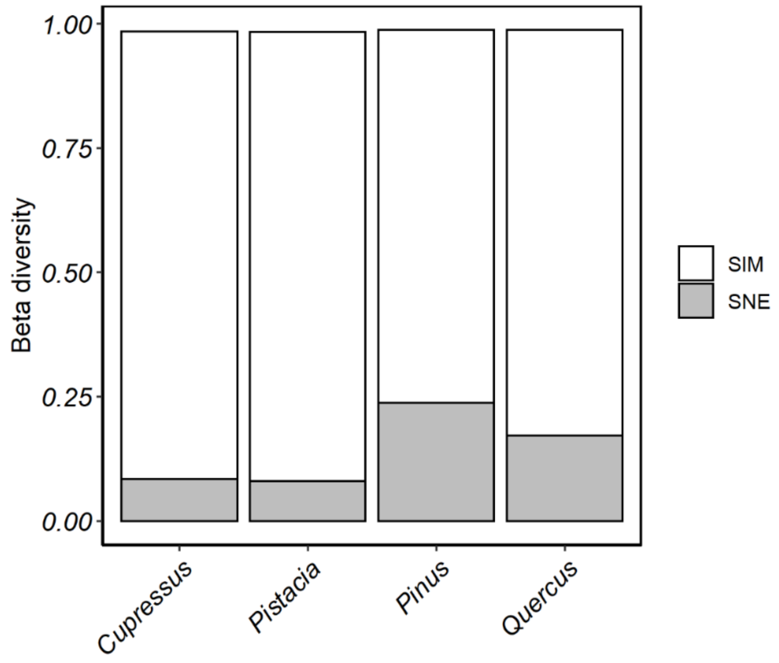


**Figure S4**. Beta-diversity estimates can be attributed to two components: spatial species turnover (white) and the nestedness of species assemblages (grey). These components were estimated using bootstrap averages of individual trees of each host tree species.

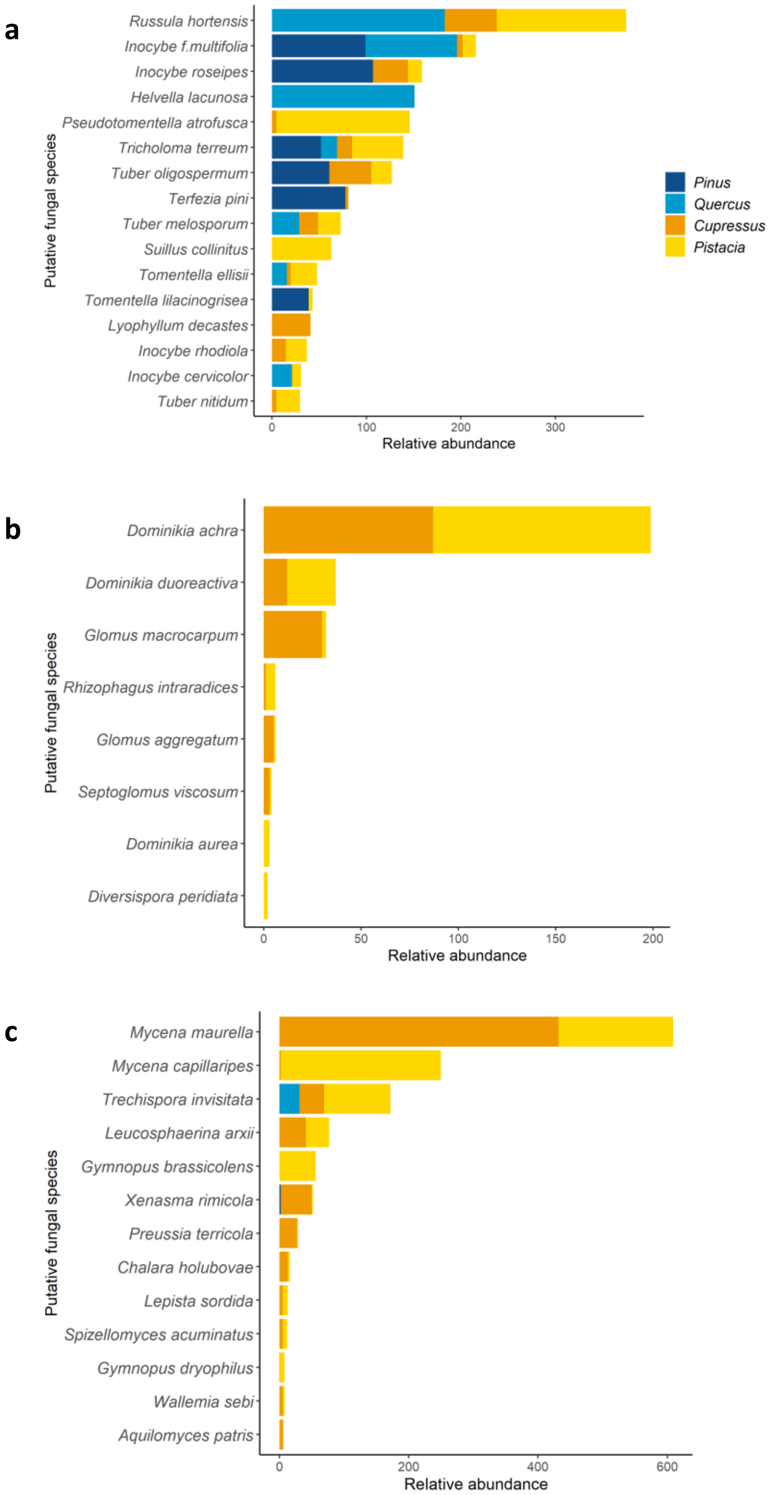


**Figure S5**. Stacked bar chart showing the relative abundance of putative fungal species: (a) EMF (b) AMF (c) SAP. Colors represent host tree species.

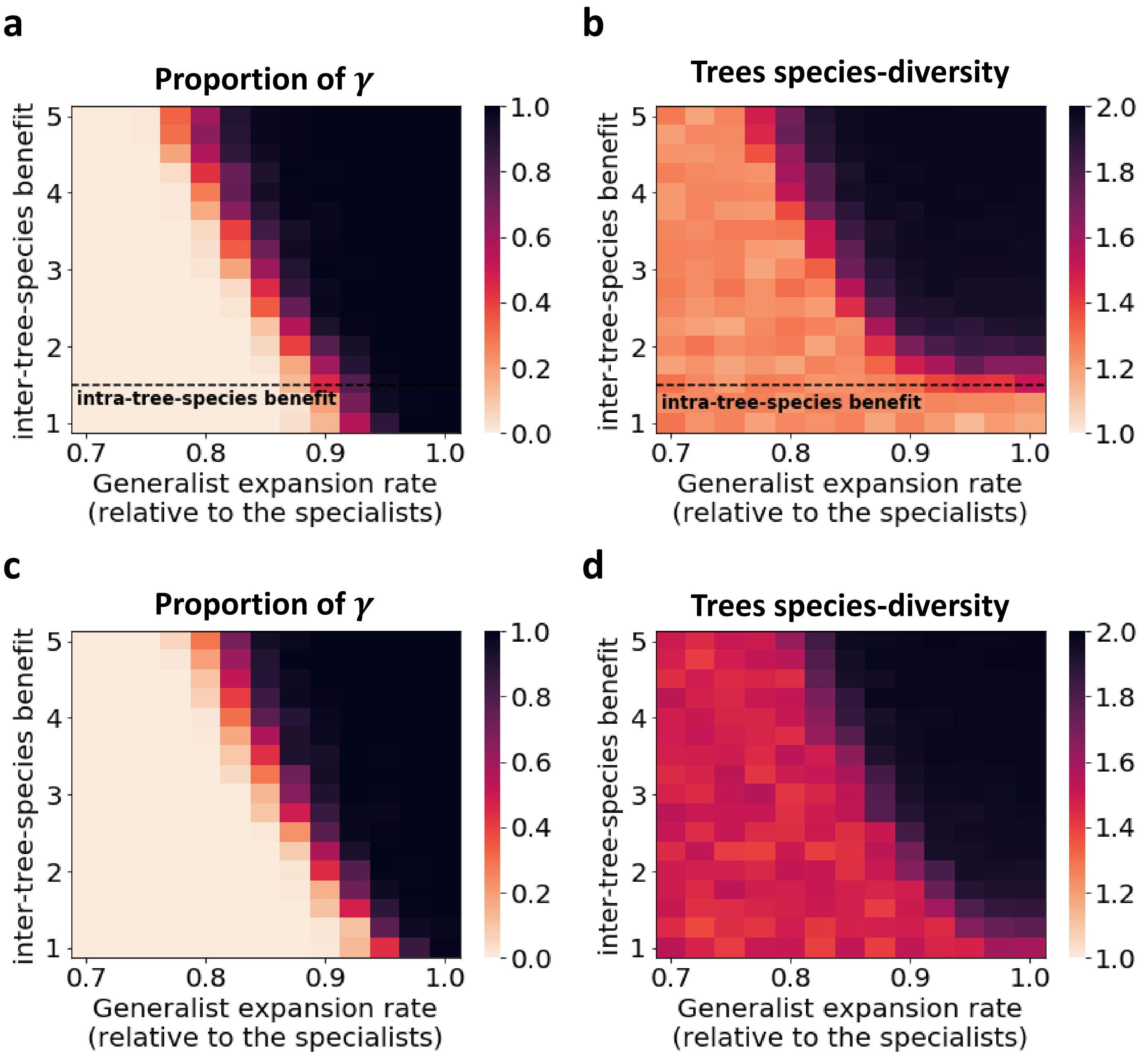


**Figure S6.** Generalist vs. specialist mycorrhiza competition models. Similarly to figures 3b and 3c in the main text, plotted are the expected proportion of $\gamma$ (a, c) and the tree-species diversity obtained by calculating $\frac{1}{x_A^2+x_B^2}$, where $x_A$ and $x_B$ are the proportions of trees $A$ and $B$, respectively (b, d). Each data point in all panels represents the average of 100 simulations. For panels (a) and (b) the simulations are stopped after 20,000 years, while in (c) and (d) after 10,000 years (similarly to figures 3b and 3c in the main text). For panels (a, b) we set the intra-tree-species benefit of mycorrhiza $\gamma$ ($b_\gamma^1$) to 1.5 (similarly to figures 3b and 3c in the main text), while for panels (c, d) it is set to 1. In all the simulations, we set the cost of cooperation to 0.2 and the expansion rate of α and β to 0.1.

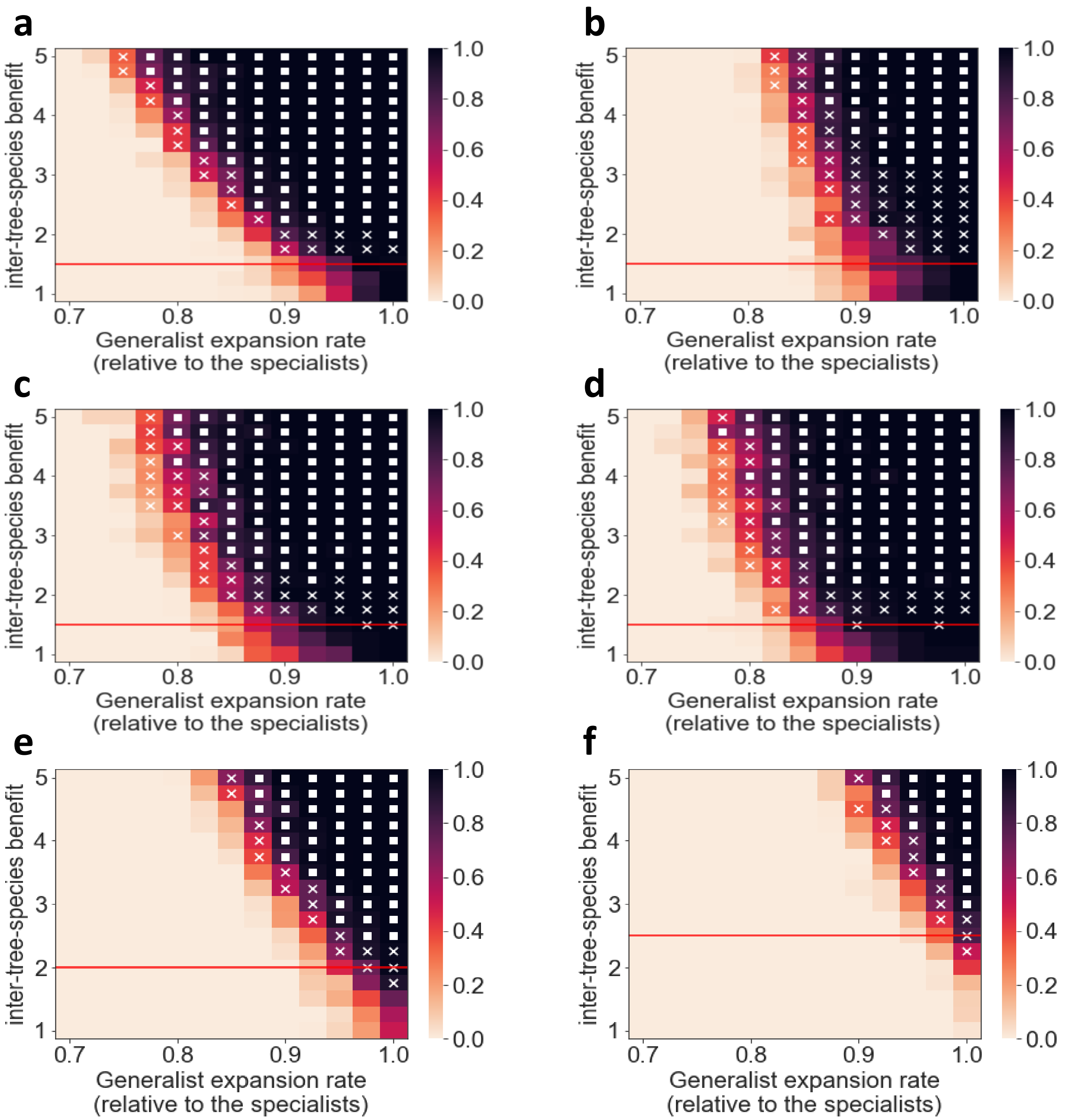


**Figure S7. Variables' robustness analysis.** The expected proportion of γ after 10,000 time points, as function of the inter-species cooperation benefit (y-axis) and of the generalist (γ) expansion rate relative to that of the specialists (α,β; x-axis). The figure show results similar to Figure 4d in the main text, but with a change in the value of one parameter in each panel. Panels (a,b) are with $c = 0.1$ and $c = 0.3$, respectively. Panels (c,d) are with specialists' expansion rate set to 0.2 and 0.3, respectively. Panels (e,f) are with intra-tree-species benefit set to 2 and 2.5, respectively.
